# Activation and Evasion of the FEAR Pathway by RNA Viruses

**DOI:** 10.1101/2024.08.22.609092

**Authors:** Emily A. Rex, Dahee Seo, Aaron Embry, Rozanne Arulanandam, Marcus M. Spinelli, Jean-Simon Diallo, Don B. Gammon

## Abstract

We recently identified the <u>F</u>ACT-<u>E</u>TS-1 <u>A</u>ntiviral <u>R</u>esponse (FEAR) pathway as an interferon-independent innate immune response that restricts DNA virus replication and is countered by poxvirus-encoded A51R proteins (Rex *et al.*, 2024, *Nature Microbiology*). The human FEAR pathway is mediated by the FACT complex, consisting of hSpt16 and SSRP1 subunits, that remodels chromatin to activate expression of the antiviral transcription factor, ETS-1. To counter this pathway, poxvirus A51R proteins tether SUMOylated hSpt16 subunits to microtubules to prevent ETS-1 expression. While these observations indicate a role for the FEAR pathway in DNA virus restriction, it was unclear if RNA viruses interact with this pathway. Here, we show that RNA viruses are also restricted by the FEAR pathway, yet encode mechanisms distinct from poxviruses to counter this response. We show vesicular stomatitis virus (VSV), a rhabdovirus, utilizes its matrix (M) protein to promote proteasome-dependent degradation of SUMOylated hSpt16 and to block ETS-1 nuclear import. Strains encoding mutant M proteins that cannot antagonize the FEAR pathway exhibit replication defects in human cells that can be rescued by hSpt16 or ETS-1 depletion. Moreover, FACT inhibitor treatment enhanced the replication of oncolytic VSV strains encoding defective M proteins in restrictive cancer cells, suggesting FEAR pathway inhibition may improve oncolytic virotherapy. Strikingly, we provide evidence that the inability of VSV M to degrade SUMOylated Spt16 in lepidopteran insect cells results in abortive infection, suggesting VSV-Spt16 interactions influence virus host range. Lastly, we show that human and murine paramyxovirus target SUMOylated Spt16 proteins for degradation in human and murine cells utilizing a conserved N-terminal motif in their accessory “C” proteins. Collectively, our study illustrates that DNA and RNA viruses have independently evolved diverse mechanisms to antagonize SUMOylated host Spt16 proteins, underscoring the physiological importance of the FEAR pathway to antiviral immunity.

## Introduction

The activation of transcription-dependent responses during infection is a fundamental aspect of eukaryotic antiviral immunity. In turn, evolutionary pressures on viruses to efficiently replicate in their eukaryotic hosts have driven the acquisition of viral countermeasures to these host responses [1–3]. Perhaps the strongest evidence for the importance of an innate immune response pathway to combating infection is the identification of pathogen-encoded antagonists of that pathway. The fact that virtually all mammalian viruses encode antagonists of the Type I interferon (IFN) response highlights the importance of this antiviral pathway [3–5]. However, relatively little is known regarding IFN-independent transcriptional response pathways that restrict virus replication and the mechanisms used by viruses to thwart such responses.

We recently reported the discovery of an IFN-independent “<u>F</u>ACT-<u>E</u>TS-1 <u>A</u>ntiviral <u>R</u>esponse (FEAR)” pathway that requires the cellular <u>Fa</u>cilitates <u>C</u>hromatin <u>T</u>ranscription (FACT) complex for activation [6]. FACT is an ancient chromatin remodeling factor that is conserved from yeast to humans. In humans, FACT is comprised of human suppressor of ty 16 homologue (hSpt16) and structure specific recognition protein 1 (SSRP1) subunits that regulate cellular gene transcription by assembling and disassembling nucleosomes at specific loci [7, 8]. Using vaccinia virus (VV), a large DNA virus belonging to the *Poxviridae* family, we showed that viral infection triggers the FACT-dependent expression of E26 transformation-specific sequence-1 (ETS-1), an antiviral transcription factor, which subsequently promotes VV restriction [6]. ETS-1 expression during infection is dependent upon FACT complexes that contain a specialized, SUMOylated form of hSpt16 (hSpt16^SUMO^)[6]. However, we found the VV-encoded A51R protein to block ETS-1 expression by outcompeting SSRP1 for direct binding to hSpt16^SUMO^ subunits in the cytosol and by tethering hSpt16^SUMO^ to microtubules [6]. VV mutant strains lacking A51R or encoding A51R mutants unable to bind hSpt16^SUMO^ strongly induce ETS-1 expression and display attenuated replication in human cell culture and in mice, suggesting the FEAR pathway is a critical component of eukaryotic antiviral immunity and that poxvirus A51R proteins function as FEAR pathway antagonists [6].

Prior to our identification of the FEAR pathway, we had serendipitously discovered that poxvirus A51R proteins could rescue the abortive infection of lepidopteran (moth and butterfly) cells by vesicular stomatitis virus (VSV), a small RNA virus belonging to the *Rhabdoviridae* family [9]. Although VSV is an arbovirus and can be naturally transmitted by blood-feeding dipteran (flies and mosquitoes) insects to mammalian hosts, lepidopterans are unnatural hosts for this virus. Thus, we suspected that A51R proteins may promote productive VSV replication in lepidopteran cells by suppressing an immune response that VSV is incapable of evading in these host cells [9]. Given that A51R proteins are only encoded by vertebrate poxviruses [9], we suspected that these viral proteins are likely immunosuppressive in insect cells because they target antiviral machinery conserved between invertebrate and vertebrate hosts. However, the nature of the host response restricting VSV and suppressed by A51R has remained elusive.

Understanding innate immune responses restricting VSV replication is important for many reasons. For one, VSV it is associated with outbreaks in livestock in the United States and thus poses a significant economic burden to the agriculture industry [10]. Moreover, the early development of a reverse genetic system for VSV [11, 12], its ability to infect a broad range of host cell types [13], and inability to cause significant disease in humans [14], have made VSV an important model for understanding fundamental aspects of nonsegmented negative-strand (NNS) RNA virus biology. VSV has also become an important vaccine platform for a wide variety of other viral diseases [14, 15].

VSV is also being investigated as an oncolytic agent in basic and clinical studies [16, 17]. A rapid and lytic replication cycle, and lack of preexisting immunity in the human population to VSV, are key advantages of its use in oncolytic virotherapy. Oncolytic VSV strains encoding either a deletion or Arg substitution at Met51 in the VSV M protein (VSV^ΔM51^ and VSV^M51R^) have been widely studied in oncolytic virotherapy [16–23]. These strains typically display attenuated replication due to their reduced ability to block host gene expression, although how wild-type VSV M normally impedes cellular gene expression is controversial [24, 25]. Prior work has shown that antiviral gene expression by the IFN response is antagonized by wild-type, but not VSV^ΔM51^ or VSV^M51R^ strains [16–19]. It has been suggested that the reduced ability of VSV^ΔM51^ and VSV^M51R^ strains to replicate in untransformed cells may be due to defective IFN antagonism [18, 21, 25, 26]. However, in transformed cells, the IFN pathway is often defective because this pathway can also activate anti-proliferative and pro-death responses that would be detrimental to cancer cell replication and survival [27]. Thus, part of the selectivity of VSV^ΔM51^ and VSV^M51R^ strains for replication in cancer cells may be in part due to defective IFN signaling in these cells. However, not all transformed cells are IFN-defective, possibly explaining why some cancer cell types are refractory to oncolytic VSV strains [18, 21, 23]. Indeed, prior work has shown that certain cancer cell lines that are refractory to VSV^ΔM51^ replication can become sensitized to infection by treatment with compounds that inhibit IFN responses [18, 21]. However, it is unclear if IFN-independent responses also contribute to VSV^ΔM51^ and VSV^M51R^ restriction in refractory cancer cell types. Thus, identifying and overcoming innate immune responses present in cancer cells that restrict oncolytic VSV strain replication is critical for improving virotherapy because enhanced VSV replication is associated with increased oncolysis and improved virotherapy outcomes [18, 20, 21, 28, 29].

Interestingly, upregulation of FACT and ETS-1 have been linked to oncogenesis and/or tumor invasion [30–32]. For example, FACT subunit upregulation is strongly associated with aggressive tumor types [31], suggesting that FACT can drive oncogenic gene expression programs that favor cancer development. This led to the development of small molecule inhibitors of FACT called “curaxins” that are currently being pursued in clinical trials for the treatment of human malignancies [33]. ETS-1 is a proto-oncoprotein and its upregulation promotes cellular transformation and tumor invasion [30, 34]. These observations suggest that the FEAR pathway involves host factors that are both antiviral and pro-tumoral. Thus, if the FEAR pathway is retained in transformed cells, it may be relevant to the restriction of oncolytic viruses and thus blocking this pathway with FACT inhibitors such as curaxins may be a logical strategy to improve the oncolytic virus replication.

With our discovery of poxvirus A51R proteins as the first DNA virus-encoded FEAR pathway antagonists [6], we wanted to use VSV as a model to investigate if RNA viruses activate and/or antagonize this pathway. Here, we show that the attenuated replication of VSV^ΔM51^/VSV^M51R^ strains is at least in part due their inability to suppress FEAR pathway activation in human cells. In contrast, VSV strains encoding wild-type M proteins effectively suppress this pathway by promoting the proteasome-dependent degradation of hSpt16^SUMO^ to abrogate ETS-1 expression and by blocking ETS-1 nuclear accumulation. We find that inhibition of the FEAR pathway in human cancer lines normally resistant to VSV^ΔM51^/VSV^M51R^ infection by hSpt16 RNA interference (RNAi) or curaxin treatment sensitizes these cancer cells to infection with these oncolytic viruses. Moreover, we show that the host range restriction of VSV in lepidopteran cells likely results from an inability of VSV M to degrade SUMOylated Spt16 proteins encoded by these insect hosts. Our data suggests that VV A51R rescues this VSV host range restriction by interacting with SUMOylated Spt16 expressed in these insect cells. Finally, we show that murine and human paramyxovirus-encoded C proteins target SUMOylated Spt16 proteins for proteasomal degradation in mouse and human cells using a conserved motif in their N-termini, suggesting that other NNS RNA viruses have independently evolved strategies to degrade these host factors. Collectively, our work illustrates that RNA viruses can also activate and antagonize the FEAR pathway and, to our knowledge, identifies VSV M and paramyxovirus C proteins as the first RNA virus-encoded FEAR pathway antagonists.

## Results

### hSpt16 or ETS-1 depletion rescues the replication defect of a VSV^M51R^ mutant

We previously showed that hSpt16 RNAi significantly enhances the replication of VSV in both A549 cells and primary neonatal human dermal fibroblasts, suggesting that FACT contributes to VSV restriction [6]. However, we wanted to ask if ETS-1 was also restrictive to VSV replication because it was possible that FACT-mediated VSV restriction is independent of ETS-1. Moreover, because VSV^ΔM51^ or VSV^M51R^ strains are thought to exhibit replication defects because of their inability to block host gene expression-dependent antiviral responses, we wondered whether these replication defects may be related to a differential ability of these mutant viruses to antagonize the FEAR pathway.

To begin to address these questions, we assessed the replication of recombinant VSV strains encoding eGFP and either wild-type (VSV-eGFP) [35] or M51R mutant M (VSV^M51R^-eGFP) proteins [35] in A549 cells transfected with small interfering RNAs (siRNAs) targeting hSpt16 or ETS-1 [6]. We found hSpt16 or ETS-1 RNAi to significantly enhance VSV-eGFP replication compared to scrambled (control) RNAi conditions, suggesting the FEAR pathway is indeed restrictive to VSV. The VSV^M51R^-eGFP strain exhibited a replication defect compared to VSV-eGFP under control RNAi conditions but strikingly, under either hSpt16 or ETS-1 RNAi conditions, this mutant replicated to titers indistinguishable from VSV-eGFP (**Fig 1A** and **B**). These results suggest that the FEAR pathway contributes to VSV restriction and that the VSV^M51R^-eGFP mutant strain is more sensitive to restriction by this pathway.

**Fig 1.**
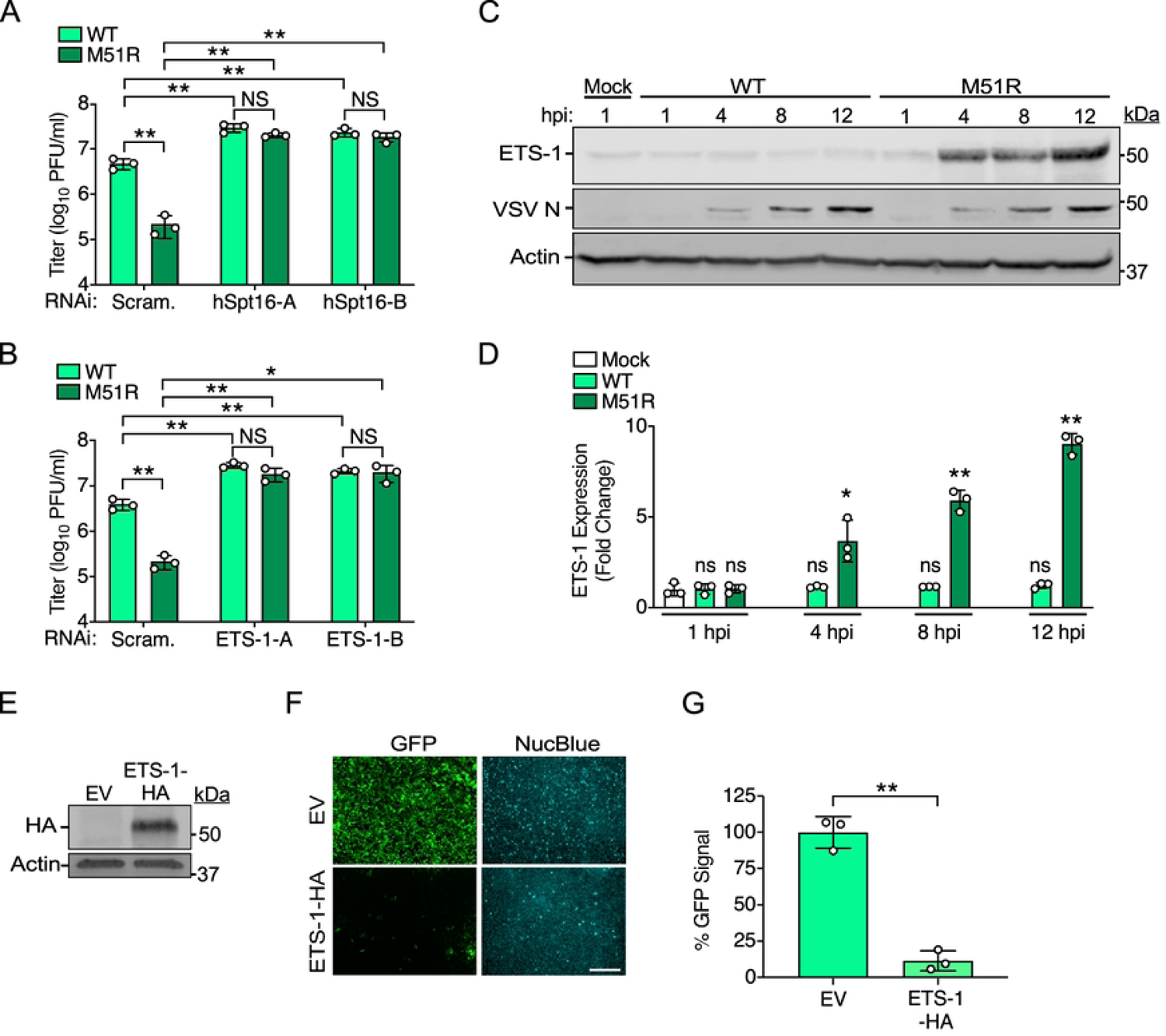
The FEAR pathway contributes to VSV restriction in human cells and is strongly activated by VSV^M51R^-eGFP. (A-B) VSV-eGFP (WT) and VSV^M51R^-eGFP (M51R) titers 24 h post-infection (hpi) (MOI=0.001) in A549 cells transfected with indicated RNAi treatments using two independent siRNAs for either hSpt16 (A) or ETS-1 (B) knockdown. Scram., scrambled siRNA. Data are means ± SD; n=3. Statistical significance in A and B were determined by unpaired two-tailed Student’s t-tests between indicated treatments. * = P< 0.05; ** = P< 0.01; ns = not significant. (C) Immunoblot (IB) of endogenous ETS-1 in A549 whole cell extract (WCE) after infection with VSV-eGFP (WT) or VSV^M51R^-eGFP (M51R) (MOI=10). (D) Densitometric quantification of ETS-1 from multiple IB experiments as in C. Data are means ± SD; n=3. Results of unpaired two-tailed Student’s t-test between ETS-1 levels in mock WCE and infected WCE are shown above each bar graph as: * = P< 0.05, ** = P< 0.01, or ns = not significant. (E) IB of U2OS cell WCE transfected with empty vector (EV) or ETS-1-HA constructs 48 h post-transfection. (F) Representative fluorescence microscopy images of U2OS cells transfected as in E and then infected with VSV-eGFP for 16 h. NucBlue was used to mark cell nuclei and to account for cell number. Scale bar = 500 μm. (G) Quantification of total GFP signal/field normalized to NucBlue signal for experiments from F. Data are means ± SD; n=3. Statistical significance was determined by unpaired two-tailed Student’s t-test between indicated treatments. ** = P< 0.01.

### Wild-type VSV, but not VSV^M51R^, inhibits ETS-1 expression to evade restriction

The results above were reminiscent of our VV studies where we found a VV strain lacking A51R (ΔA51R) to exhibit a replication defect that could be rescued by FACT subunit or ETS-1 depletion. We also found this ΔA51R strain to more strongly induce ETS-1 expression than wild-type VV [6]. Thus, we next used infection time courses and immunoblots (IBs) to determine if ETS-1 levels differ between VSV-eGFP and VSV^M51R^-eGFP infections. Interestingly, while ETS-1 protein levels were not significantly different between mock- and VSV-eGFP-infected cells, VSV^M51R^-eGFP-infected WCE showed significantly elevated ETS-1 by 4 h post-infection (hpi) (**Fig 1C** and **D**). These results indicate that infection with the VSV^M51R^-eGFP mutant induces ETS-1 expression while wild-type (VSV-eGFP) infection does not induce ETS-1.

We hypothesized that wild-type VSV suppresses ETS-1 expression to prevent activation of antiviral gene expression programs that require elevated ETS-1 levels. Therefore, we next asked if ETS-1 overexpression prior to infection could suppress VSV-eGFP infection. To do this, we overexpressed an HA-tagged human ETS-1 construct (ETS-1-HA) in cells for 48 h (**Fig 1E**) and then challenged cells with VSV-eGFP. Using fluorescence microscopy, we quantified GFP signals as a readout for infection and normalized these signals to total cell number by labeling cell nuclei with NucBlue. Compared to cultures transfected with empty vector (control), ETS-1-HA-transfected cells displayed significantly reduced GFP signal (**Fig 1F** and **G**). Collectively, these data suggest ETS-1 expression results in a block to VSV infection but VSV-eGFP evades this restriction by suppressing ETS-1 expression. In contrast, VSV^M51R^-eGFP is unable to block ETS-1 induction.

### VSV^M51R^ infection induces ETS-1 expression independently of RIG-I and the IFN response

We showed ΔA51R VV strains to induce ETS-1 expression in the absence of an intact IFN response [6]. Therefore, we wanted to determine if ETS-1 induction by VSV^M51R^-eGFP infection was also independent of the IFN response. Since the pattern recognition receptor RIG-I is required for the activation of IFN responses during VSV infection of non-immune cell types [36], we first examined ETS-1 induction in RIG-I-deficient A549 cells. No significant differences in ETS-1 induction were found between wild-type and RIG-I knockout cells during VSV^M51R^-eGFP infection (**Fig S1A and B**). We also quantified ETS-1 expression levels in A549 cells lacking IRF3 or STAT1, two critical components of the IFN signaling pathway [37, 38]. ETS-1 induction by VSV^M51R^-eGFP infection was unaffected in these IFN pathway knockout cell lines (**Fig S1C-F**). Collectively, these data indicate that, like VV, VSV activates ETS-1 expression independently of the IFN response.

### VSV, but not VSV^M51R^, promotes hSpt16^SUMO^ depletion to prevent ETS-1 induction

During VV infection, FACT complexes comprised of hSpt16^SUMO^ and SSRP1 subunits are required to induce ETS-1 expression [6]. Therefore, we hypothesized that VSV-eGFP may suppress ETS-1 expression by altering FACT subunit levels. Using infection time courses and IBs, we determined if hSpt16^SUMO^ or SSRP1 levels differ between VSV-eGFP and VSV^M51R^-eGFP infections. It is important to note that both hSpt16^SUMO^ and SUMOless hSpt16 forms can be resolved from one another by SDS-PAGE using low percentage acrylamide protein gels and then simultaneously detected via IB with hSpt16 antibodies (Ab) [6]. Strikingly, our IBs revealed a clear and significant reduction of hSpt16^SUMO^ (upper band in **Fig 2A**) levels compared to mock-infected WCE during VSV-eGFP infection by 4 hpi and hSpt16^SUMO^ became virtually undetectable by 8 hpi. In contrast, hSpt16^SUMO^ levels remained unchanged throughout the entire VSV^M51R^-eGFP infection time course (**Fig 2A** and **B**). Interestingly, SUMOless hSpt16 (lower band in **Fig 2A**) or SSRP1 levels did not change during infection with either virus (**Fig 2C** and **D**), suggesting that VSV infection leads to the specific depletion of hSpt16^SUMO^ subunits.

**Fig 2.**
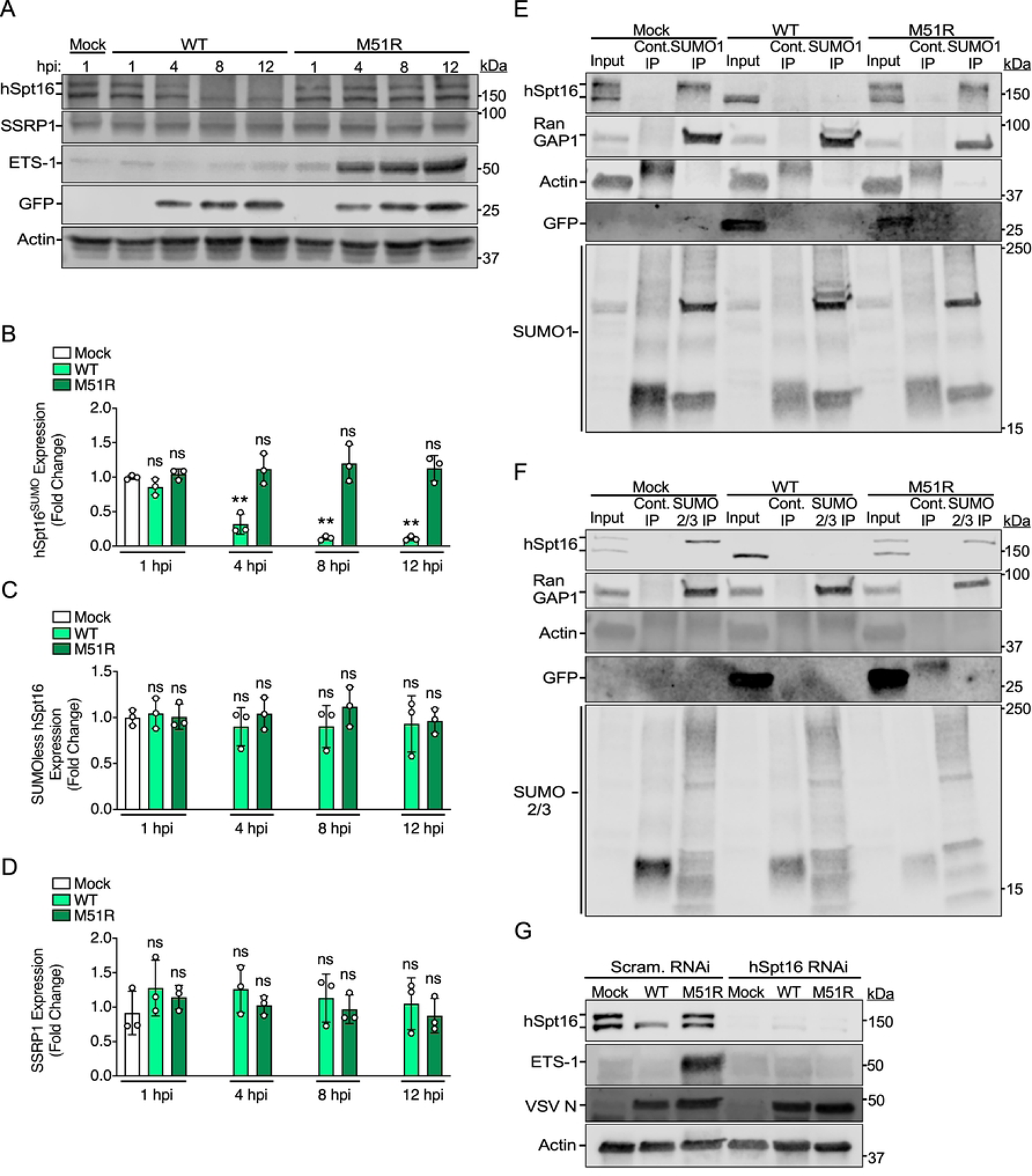
VSV, but not VSV^M51R^, promotes hSpt16^SUMO^ depletion to prevent ETS-1 induction. (A) IB of FACT subunit levels in WCE from A549 cells infected with VSV-eGFP (WT) or VSV^M51R^-eGFP (M51R) for indicated hpi (MOI=10). (B-D) Densitometric quantification of IB experiments in A for hSpt16^SUMO^ (B) SUMOless hSpt16 (C) or SSRP1 levels (D). Data are means ± SD; n=3. In B-D, results of unpaired two-tailed Student’s t-test between protein levels in mock WCE and infected WCE are shown above each bar graph as: * = P< 0.05, ** = P< 0.01, or ns = not significant. (E-F) IB of immunoprecipitated SUMO-1-(E) or SUMO-2/3-(F) conjugated protein fractions in WCE from mock-, VSV-eGFP -, or VSV^M51R^-eGFP-infected A549 cells (MOI=3) 18 hpi. Cont., control IP. RanGAP1 is a known SUMOylated protein and used as a control for enrichment of SUMOylated fractions [39]. GFP is a marker for infection. (G) IB of endogenous ETS-1 and hSpt16 levels and in WCE of scrambled- or hSpt16-siRNA-treated A549 cells 8 hpi with indicated strains (MOI=10).

When comparing the effect of MOI on hSpt16^SUMO^ depletion during VSV-eGFP infection, we noticed that complete depletion of hSpt16^SUMO^ levels occurred earlier (by 4 hpi) when using higher MOIs (e.g. 10-30) versus during lower MOI (e.g. 3) infections. However, the VSV^M51R^-eGFP strain could not deplete hSpt16^SUMO^ at any MOI or time point (**Fig 2A and S2)**.

To confirm these data, and to determine if loss of hSpt16^SUMO^ levels during VSV-eGFP infection was not due to a global down regulation of intracellular SUMOylation, we used Ab against SUMO-1 or SUMO-2/3 to immunoprecipitate (IP) SUMOylated protein fractions from mock-, VSV-eGFP-, or VSV^M51R^-eGFP-infected WCE [6]. As expected, hSpt16^SUMO^ was specifically depleted in VSV-eGFP-infected WCE while present in VSV^M51R^-eGFP-infected lysates. Moreover, no overt reductions in either total SUMOylation banding patterns or levels of SUMOylated-RanGAP1 (an abundant, SUMOylated protein [39]) were observed in VSV-eGFP versus VSV^M51R^-eGFP infections (**Fig 2E** and **F**). These data suggest that VSV-eGFP can promote depletion of hSpt16 that is SUMOylated with either SUMO-1 or SUMO-2/3 and that the specific loss of hSpt16^SUMO^ levels during VSV-eGFP infection is not due to a generalized decrease in intracellular SUMOylation.

Finally, because hSpt16^SUMO^ is required for ETS-1 induction during ΔA51R VV infection [6], we wanted to determine if hSpt16 RNAi treatment would inhibit ETS-1 induction by VSV^M51R^-eGFP infection. Consistent with our VV studies, hSpt16 RNAi treatment abrogated ETS-1 induction after VSV^M51R^-eGFP infection (**Fig 2G**). Collectively, these data indicate that the inability of VSV^M51R^-eGFP to deplete hSpt16^SUMO^ results in robust ETS-1 induction during infection.

### VSV M is sufficient to promote proteasomal degradation of cytoplasmic hSpt16^SUMO^ and to block ETS-1 expression

Given that the only difference between VSV-eGFP and VSV^M51R^-eGFP strains is the M51R substitution in the VSV M protein, we hypothesized that this viral factor was responsible for hSpt16^SUMO^ depletion. Consistent with this, a VSV^ΔM51^ strain expressing GFP (VSV^ΔM51^-GFP [18]) was also unable to deplete hSpt16^SUMO^ levels when compared to its wild-type VSV-GFP control strain (**Fig S3**). However, we still conducted a screen of 4 of the 5 VSV-encoded proteins to determine if any of these viral factors could reduce hSpt16^SUMO^ levels when expressed from plasmids in uninfected human cells. The VSV L protein, encoding the viral RNA polymerase, was excluded from this screen as it seemed unlikely to be involved in immune evasion. Consistent with our hypothesis, only expression of Flag-tagged VSV M (VSV M-Flag) resulted in hSpt16^SUMO^ depletion (**Fig 3A**). Moreover, expression of either VSV M^ΔM51^-Flag or VSV M^M51R^-Flag mutant constructs failed to deplete hSpt16^SUMO^ levels (**Fig 3B**), consistent with our infection results. These data indicate that wild-type VSV M proteins, but not M51R or ΔM51 mutants, are sufficient for hSpt16^SUMO^ depletion in the absence of other VSV proteins.

**Fig 3.**
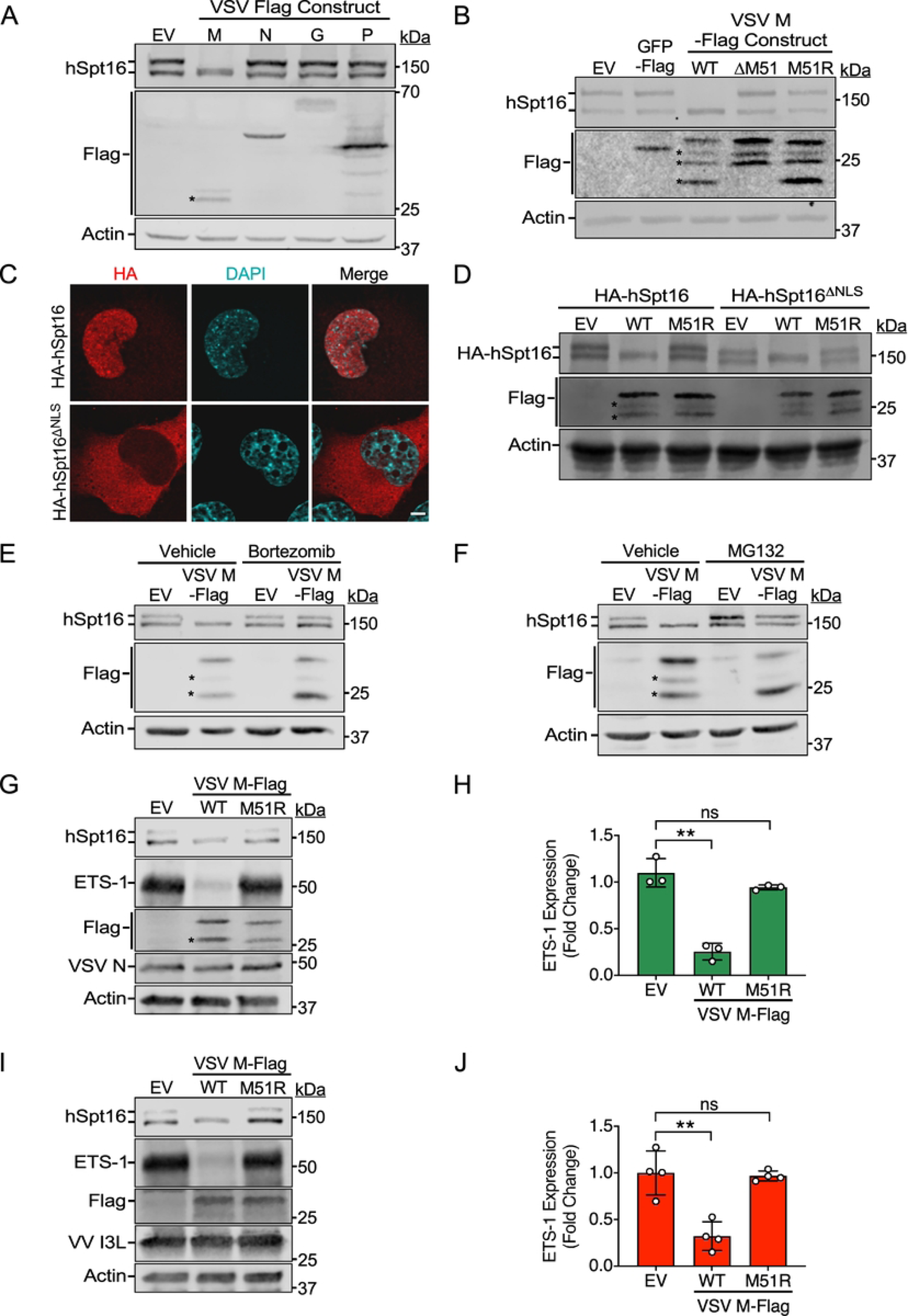
VSV M is sufficient to promote proteasomal degradation of cytoplasmic hSpt16^SUMO^ and to block ETS-1 expression. (A) IB of endogenous hSpt16 in 293T WCE transfected with indicated Flag-tagged VSV protein expression constructs for 24 h. Empty vector, EV. Asterisks indicate probable VSV M degradation products. (B) IB of endogenous hSpt16 in 293T WCE transfected with indicated VSV M expression constructs 24 h post-transfection. EV and Flag-GFP constructs were negative controls. Asterisks indicate probable VSV M degradation products. (C) Immunofluorescence (IF) images of U2OS cells transfected with indicated HA-tagged hSpt16 constructs for 24 h. Scale bar = 5 μm. (D) IB of 293T WCE 24 h post-transfection with indicated HA-tagged hSpt16 constructs along with either EV, VSV M-Flag (WT), or VSV M^M51R^-Flag (M51R) pcDNA3 vectors. Asterisks indicate probable VSV M degradation products. (E-F) 293T WCE transfected with indicated VSV M expression constructs in the absence or presence of bortezomib (E) or MG132 (F) proteasome inhibitor treatment. Vehicle (DMSO) treatments served as negative controls. Asterisks indicate probable VSV M degradation products. (G) IB of endogenous ETS-1 in U2OS WCE after transfection with indicated expression constructs for 24 h and then infected with VSV^M51R^-eGFP (MOI =10) for 8 h. (H) Densitometric quantification of IB experiments in G. Data are means ± SD; n=3. (I) IB of endogenous ETS-1 in U2OS WCE after transfection with indicated expression constructs for 24 h and then infected with ΔA51R VV strain (MOI =10) for 8h. (J) Densitometric quantification of IB experiments in I. Data are means ± SD; n=4. Statistical significance in H and J were determined by unpaired two-tailed Student’s t-tests between indicated treatments. ** = P< 0.01; ns = not significant.

VSV M can inhibit host gene expression by binding to Rae1, a component of the cellular Rae1-Nup98 export complex that promotes mRNA export from the nucleus [25]. However, the mechanism by which VSV M-Rae1 interaction inhibits host gene expression is controversial [24, 25]. In one proposed model, VSV M binds Rae1 to inhibit Rae1-Nup98-mediated mRNA export from the nucleus while an alternative model suggests VSV M may usurp Rae1-Nup98 complexes to inhibit cellular gene transcription [24, 25]. Notably, VSV M51R/ΔM51 M mutants cannot interact with Rae1 [25]. Despite the unresolved nature of VSV M-Rae1 interaction models, we wanted to determine if Rae1 or Nup98 was involved in hSpt16^SUMO^ level regulation. Thus, we tested if depletion of either Rae1 or Nup98 altered hSpt16^SUMO^ levels in mock-infected cells or inhibited hSpt16^SUMO^ depletion during VSV-eGFP infection. In either case, we observed no effect of Rae1 or Nup98 depletion on hSpt16^SUMO^ levels (**Fig S4**). This suggests that VSV M antagonism of the FEAR pathway cannot be explained by its interaction with Rae1-Nup98 complexes.

We next wanted to gain additional insight into the mechanism by which VSV M depletes hSpt16^SUMO^ levels. Given that VV A51R blocks the FEAR pathway by binding to cytosolic hSpt16^SUMO^ subunits, we next asked if VSV M also targets cytosolic hSpt16^SUMO^ subunits. To do this, we co-expressed VSV M-Flag constructs with either a wild-type or nuclear localization sequence (NLS)-deleted (NLS) HA-tagged hSpt16 constructs we previously generated [6]. Importantly, we confirmed that wild-type HA-hSpt16 displayed both cytosolic and nuclear localization while the HA-hSpt16^ΔNLS^ mutant was restricted to the cytosol (**Fig 3C**). Upon co-transfection with wild-type VSV M-Flag, the SUMOylated forms of both wild-type and NLS-deleted HA-hSpt16 constructs were depleted (**Fig 3D**), indicating VSV M-Flag likely targets hSpt16^SUMO^ subunits in the cytoplasm. To determine if VSV M promotes proteasome-dependent degradation of hSpt16^SUMO^, we expressed VSV M-Flag in human cells in the absence or presence of proteasome inhibitors bortezomib or MG132 and assessed hSpt16^SUMO^ levels by IB. Interestingly, hSpt16^SUMO^ depletion was blocked in the presence of either proteasome inhibitor (**Fig 3E** and **F**), suggesting VSV M promotes the proteasome-dependent degradation of hSpt16^SUMO^.

Finally, we asked if plasmid-based expression of VSV M-Flag was sufficient to block ETS-1 induction by either VSV^M51R^-eGFP (**Fig 3G** and **H**) or ΔA51R VV (**Fig 3I** and **J**) infection. In both cases prior expression of VSV M-Flag, but not VSV M^M51R^-Flag, was able to suppress ETS-1 induction by either of these mutant viruses (**Fig 3G-J**). The ability of VSV M to complement the ΔA51R strains inability to block ETS-1 expression demonstrates VSV M is a bona fide FEAR pathway antagonist that can function independently of other VSV proteins. Collectively, our prior and current findings suggest that VV and VSV have independently evolved unique strategies to inhibit FACT-dependent ETS-1 expression by either tethering hSpt16^SUMO^ to microtubules (VV A51R) or by promoting hSpt16^SUMO^ degradation (VSV M), illustrating that FEAR pathway suppression is important for both DNA and RNA viruses.

### VSV M encodes a second mechanism to inhibit ETS-1 function: blockade of nuclear import

While the specific antiviral function that ETS-1 performs after FEAR pathway activation is still an active area of investigation, ETS-1 is known to regulate host transcription in the nucleus. Thus, we determined if ETS-1 localizes to the nucleus during VSV infection. First, we used immunofluorescence microscopy to examine endogenous ETS-1 localization in mock-, VSV-eGFP-, or VSV^M51R^-eGFP-infected human A549 cells. In mock-infected cells, both cytosolic and nuclear ETS-1 staining was observed, consistent with the known nucleocytoplasmic cycling of ETS-1 [40, 41]. However, during VSV-eGFP infection, ETS-1, was largely excluded from the nucleus and restricted to the cytosol, suggesting that VSV infection impairs ETS-1 nuclear translocation. In contrast, ETS-1 signals were stronger and predominantly nuclear in VSV^M51R^-eGFP-infected cells (**Fig 4A**). To complement our microscopy experiments, we conducted cell fractionation and IB studies to quantify the percentage of total ETS-1 protein in nuclear fractions. Consistent with our microscopy results, ∼37% of total ETS-1 was found in nuclear fractions of mock-infected cells, while only ∼10% of ETS-1 was nuclear during VSV-eGFP infection. In contrast, ∼78% of ETS-1 was found in the nucleus during VSV^M51R^-eGFP infection (**Fig 4B** and **C**). Importantly, immunofluorescence experiments in U2OS cells also demonstrated a similar exclusion of ETS-1 from the nucleus during VSV-eGFP infections as observed in A549 cells (**Fig 4D**), suggesting this was not cell type-specific. Finally, to determine if VSV M was able to impede ETS-1 nuclear localization in the absence of other VSV proteins, we co-transfected the ETS-1-HA expression vector with empty vector, VSV M-Flag, or VSV^M51R^-Flag constructs into U2OS cells. In the presence of empty vector or VSV^M51R^-Flag constructs, ETS-1-HA displayed both nuclear and cytosolic localization. However, in the presence of wild-type VSV M-Flag, ETS-1-HA staining was largely excluded from the nucleus and restricted to the cytoplasm (**Fig 4E**).

**Fig 4.**
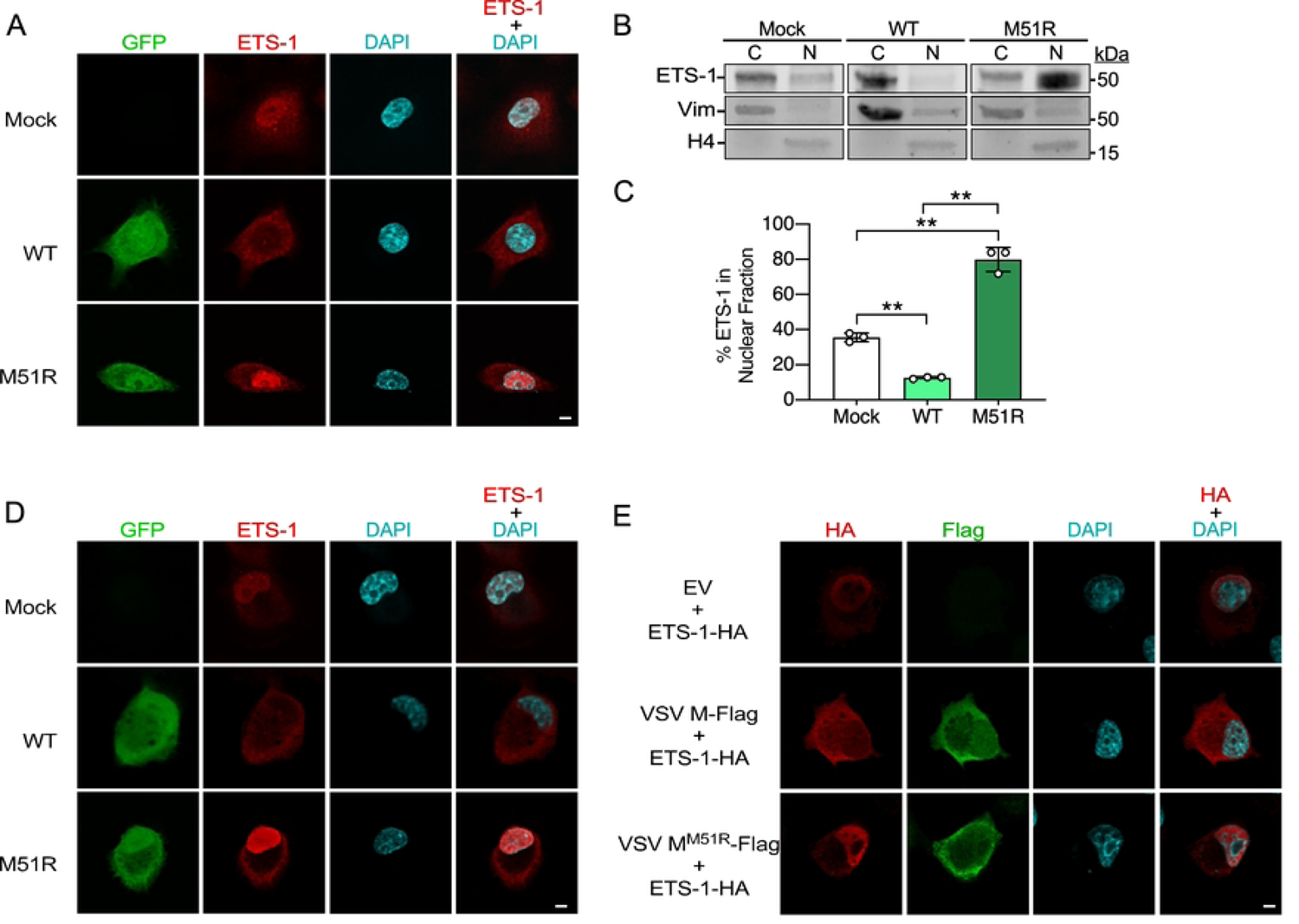
VSV M inhibits ETS-1 nuclear accumulation. (A) IF staining of ETS-1 in mock-, VSV-eGFP (WT)-, or VSV^M51R^-eGFP (M51R)-infected A549 cells 12 hpi (MOI=10). (B) IB of A549 cytoplasmic (C) and nuclear (N) fractions 12 hpi with indicated strains (MOI=10). Vimentin marks cytosolic fractions while histone H4 marks nuclear fractions [6]. (C) Densitometric quantification of multiple IB experiments as in B expressed as percentage of total (cytosolic+nuclear fractions) ETS-1 protein present in nuclear fractions. Data are means ± SD; n=3. Statistical significance was determined by unpaired two-tailed Student’s t-test between indicated treatments. ** = P< 0.01. (D) IF staining of ETS-1 in mock-, VSV-eGFP-, or VSV^M51R^-eGFP-infected U2OS cells 12 hpi (MOI=10). (E) IF staining of U2OS cells 24 h post co-transfection with ETS-1-HA expression vectors and either EV, VSV M-Flag, or VSV M^M51R^-Flag expression vectors. Scale bars in A, D, and E = 5 μm.

Collectively, these data indicate that, during infection, ETS-1 accumulates in the nucleus in the presence of the M51R mutant M protein but not in the presence of wild-type VSV M. Moreover, our transfection experiments suggest VSV M can prevent ETS-1 nuclear accumulation in the absence of other VSV proteins, indicating it encodes an additional mechanism of antagonizing the FEAR pathway-blocking ETS-1 nuclear import.

### FEAR pathway inhibition enhances oncolytic VSV strain replication in refractory cancer cells

VSV^ΔM51^ and VSV^M51R^ viruses are being pursued for use in oncolytic virotherapy. While effective at infecting some human cancer cell types, some transformed cell types have proven to be refractory to infection by these oncolytic VSV strains, posing a major challenge to their wide-spread use [18, 21, 23]. Thus, we asked if the FEAR pathway may contribute to restriction of oncolytic VSV strain replication in human cancer lines known to be refractory to VSV^ΔM51^/VSV^M51R^ viruses. First, we measured replication of VSV^ΔM51^-GFP in human 786-0 renal adenocarcinoma cells, which are relatively resistant to infection with this virus [18, 21], after hSpt16 RNAi. As shown in **Fig 5A**, VSV^ΔM51^-GFP replication was significantly increased in 786-0 cells after hSpt16 depletion by RNAi. Importantly, we confirmed that both hSpt16 and ETS-1 were reduced in 786-0 cells infected with VSV^ΔM51^-GFP during hSpt16 knockdown (**Fig 5B**). These data suggest the FEAR pathway contributes to the restriction of this virus in this restrictive cell type.

**Fig 5.**
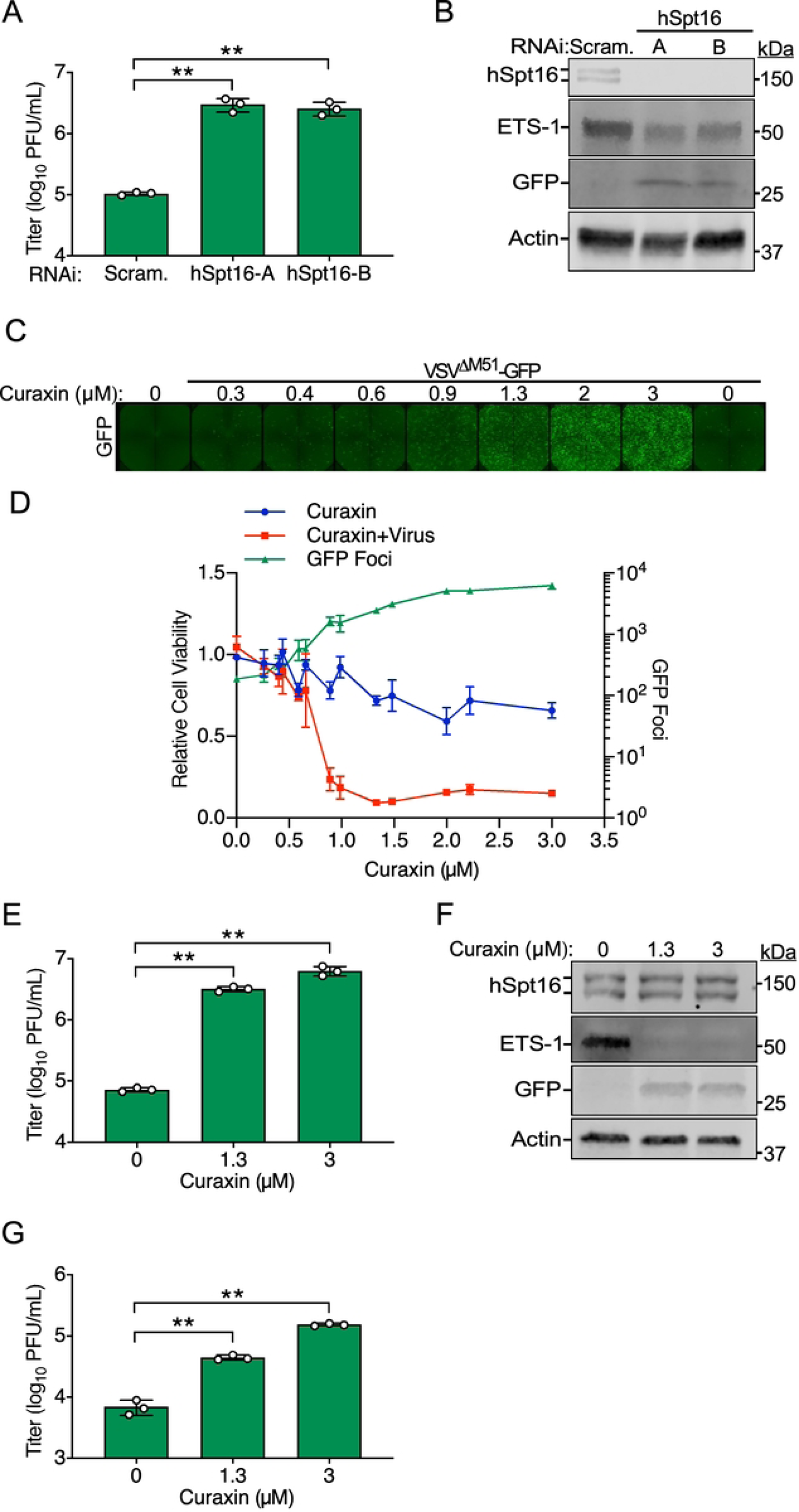
Suppression of FEAR pathway activation promotes oncolytic VSV strain replication in refractory cancer cell lines. (A) VSV^ΔM51^-GFP titers 24 hpi (MOI=0.01) in 786-0 cells transfected with indicated RNAi treatments. Data are means ± SD; n=3. (B) IB of 786-0 WCE from experiments in A at 24 hpi. (C) Representative fluorescence microscopy images 24 hpi with VSV^ΔM51^-GFP (MOI=0.05) in 786-0 cells in the presence of indicated doses of CBL0137. (D) Quantification of GFP foci 24 hpi (green line, read on right y-axis) after VSV^ΔM51^-GFP infection of 786-0 cells in the absence or presence of indicated doses of CBL0137. Cell viability 48 h post-treatment (read on left y-axis) of cultures containing only CBL0137 (blue line) or CBL0137+ VSV^ΔM51^-GFP (red line) is also shown. Data are means ± SD; n=3. (E) Titer of VSV^ΔM51^-GFP-infected 786-0 cells 24 hpi (MOI=0.01) in the presence of indicated doses of CBL0137. Data are means ± SD; n=3. (F) IB of 786-0 WCE from experiments in E at 24 hpi. (G) Titer of VSV^ΔM51^-GFP-infected RPMI 7951 cells 24 hpi (MOI=0.01) in the presence of indicated doses of CBL0137. Data are means ± SD; n=3. Statistical significance in A, E, G, and H were determined by unpaired two-tailed Student’s t-test between indicated treatments. * = P< 0.05; ** = P< 0.01; ns, not significant.

Since the use of hSpt16 RNAi as a therapeutic strategy to inhibit the FEAR pathway is likely to be technically challenging *in vivo*, we conducted a “proof of principle” series of cell culture experiments to determine if treatment with the small molecule FACT inhibitor, CBL0137, could similarly enhance VSV^ΔM51^-GFP replication in restrictive cancer cell types. CBL0137 is a second-generation curaxin and has broad activity against diverse cancer cell types in culture and in mouse models and is being pursued in clinical trials as a therapeutic for various malignancies [33, 42–45]. Curaxins inhibit FACT by causing this complex to become trapped on chromatin, preventing it from functioning properly in gene expression regulation [33]. First, we used quantitative fluorescence microscopy and GFP signal as a simple readout of VSV^ΔM51^-GFP replication in 786-0 cells in the presence of a broad range of CBL0137 concentrations 24 hpi. Addition of CBL0137 to media 1 hpi led to a dose-dependent increase in GFP signal, peaking at an ∼34-fold increase in GFP signal with addition of 3 μM CBL0137 compared to infected cultures without curaxin (**Fig 5C** and **D, right axis**). We also analyzed cell viability via fluorometric resazurin assays of parallel 786-0 cultures treated with either CBL0137 alone or VSV^ΔM51^-GFP+CBL0137 after 48 h and found a clear synergistic effect of the combined treatment on cancer cell killing. For example, in cultures treated with 1.3 μM CBL0137 only, cell viability was ∼72% of untreated cultures. However, combining 1.3 μM curaxin with VSV^ΔM51^-GFP treatment reduced 786-0 cell viability to ∼10% of controls (**Fig 5D, left axis**). Importantly, we verified that the increased GFP signal observed in CBL0137-treated cultures correlated with elevated VSV^ΔM51^-GFP titers (**Fig 5E**) and with inhibition of ETS-1 expression (**Fig 5F**). We also tested the human melanoma cell line, RPMI 7951, which is resistant to infection with VSV^M51R^ oncolytic strains [23] and observed enhanced VSV^ΔM51^-GFP replication in these cells in the presence of CBL0137 (**Fig 5G**), suggesting curaxin may sensitize other types of cancer cells to oncolytic VSV strain infection. Collectively, these data suggest that hSpt16 depletion or FACT inhibition with curaxins can enhance the replication of an oncolytic VSV strain lacking a functional FEAR pathway antagonist in restrictive human cancer cell types.

### FACT contributes to VSV host range restriction in *Lymantria dispar* insect cells

We originally became interested in understanding poxvirus A51R protein function after discovering that these proteins rescue the post-entry abortive infection of VSV in *Lymantria dispar* (gypsy moth or spongy moth)-derived LD652 cells [9]. With the knowledge that VV A51R binds to hSpt16^SUMO^ to antagonize its antiviral function [6] and that Spt16 proteins are conserved in all eukaryotes, we wondered if VV A51R-mediated rescue of VSV in LD652 cells related to its ability to antagonize insect Spt16 function.

After sequencing the *L. dispar* genome [46], we discovered that hSpt16 and *L. dispar* Spt16 (LdSpt16) proteins share an overall a.a. identity of ∼60% [6]. However, alignment of the A51R-binding domain in hSpt16 [6] with the corresponding region in LdSpt16 showed this region to be ∼73% identical between these Spt16 proteins (**Fig S5**). However, because VV A51R only binds SUMOylated forms of hSpt16, it was important to determine if LdSpt16 was also SUMOylated. Interestingly, IBs of LD652 WCE revealed two LdSpt16 forms as found with hSpt16 in human cells. Treatment of LD652 cultures with tannic acid, a global SUMOylation inhibitor [47], eliminated the upper LdSpt16 form on IBs, suggesting this band represents SUMOylated LdSpt16 (LdSpt16^SUMO^) (**Fig 6A**). To determine if VV A51R interacts with LdSpt16^SUMO^, we infected LD652 cells with a recombinant VV strain encoding Flag-tagged A51R (FA51R) under its natural *A51R* gene promoter (ΔA51R^FA51R^) and then performed co-immunoprecipitation [6]. Interestingly, LdSpt16^SUMO^, but not SUMOless LdSpt16 proteins, specifically co-immunoprecipitated with FA51R in LD652 WCE (**Fig 6B)**, suggesting VV A51R can interact with SUMOylated Spt16 proteins in this insect host. Next, we tested if A51R-LdSpt16 interactions were required for A51R-mediated rescue of VSV replication in LD652 cells. To do this, we transfected LD652 cells with a control expression vector encoding Flag-tagged GFP (FGFP) or vectors encoding wild-type FA51R, or a FA51R mutant that cannot bind to SUMOylated Spt16 proteins FA51R^158-162Ala^ [6]. 48 h post-transfection, we challenged cells with a recombinant VSV strain encoding firefly luciferase (VSV-LUC) [48]. We then assessed VSV-LUC gene expression with luciferase assays and measured viral titers in collected LD652 culture supernatants [9]. While the wild-type FA51R construct was able to rescue both VSV gene expression and replication, the FA51R^158-162Ala^ mutant was completely defective in this regard (**Fig 6C** and **D**). These results suggest that VV A51R-mediated rescue of VSV requires A51R-LdSpt16^SUMO^ interaction.

**Fig 6.**
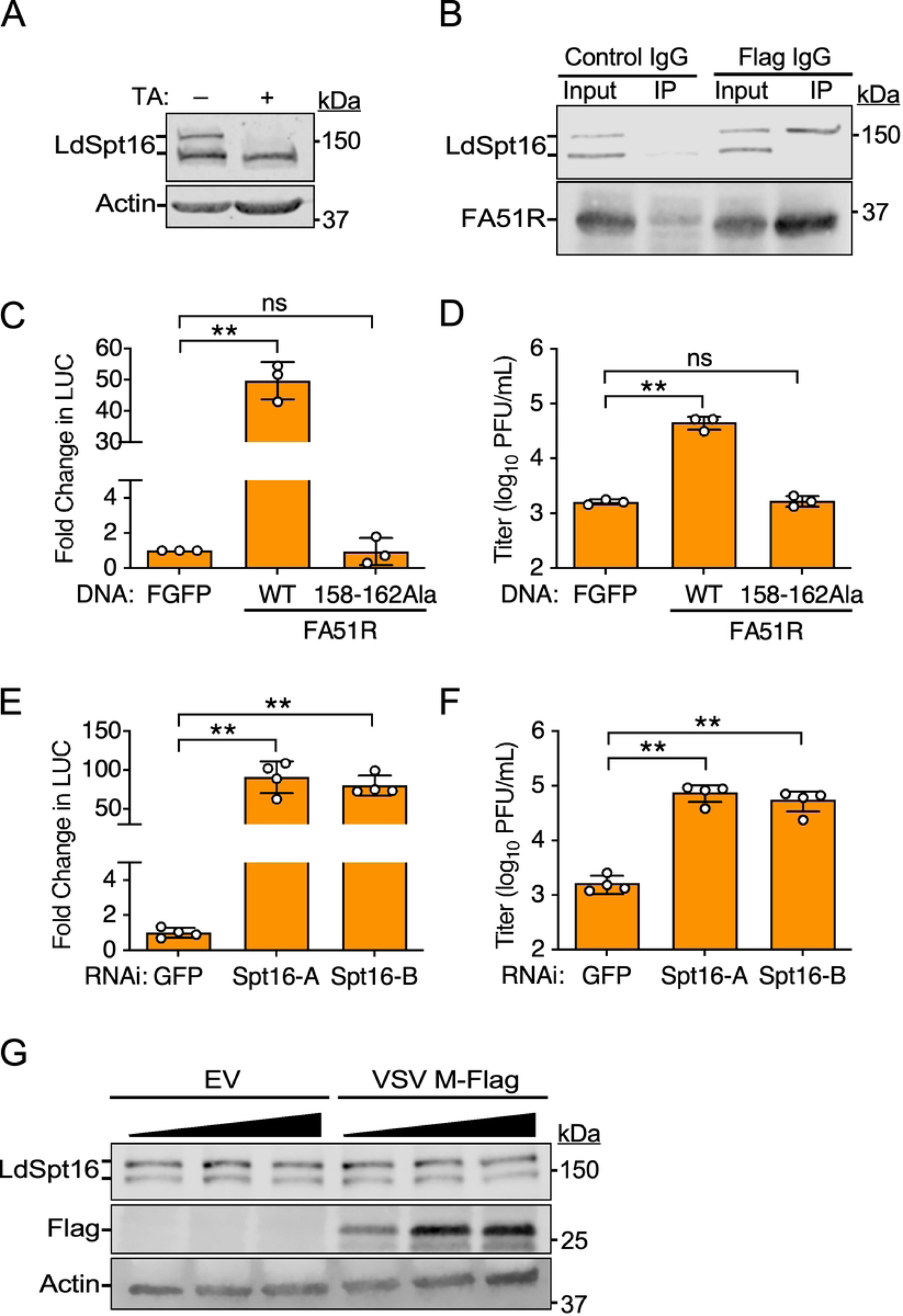
Abortive VSV replication in *L. dispar* cells can by complemented by VV A51R expression or by Spt16 RNAi. (A) IB of *L. dispar* Spt16 (LdSpt16) in LD652 WCE in the absence or presence of tannic acid (TA) treatment. (B) Co-IP of endogenous LdSpt16 with FA51R in ΔA51R^FA51R^-infected (MOI=25) LD652 WCE. (C) Fold-change in luciferase (LUC) signal in LD652 cells transfected with indicated FA51R expression constructs for 24 h and then infected with VSV-LUC (MOI=10) for 48 h. Fold-change in LUC is plotted as relative to LUC signal in Flag-GFP (FGFP) transfection treatments (negative control). Data are means ± SD; n=3. (D) VSV-LUC titers from supernatants collected from C. Data are means ± SD; n=3. (E) Fold-change in LUC signal after 72 h of transfection with dsRNAs (for RNAi) targeting either GFP (control) or LdSpt16 (Spt16-A/B) and infection with VSV-LUC (MOI=10) for 48 h. Data are means ± SD; n=4. (F) VSV-LUC titers from supernatants collected from E. Data are means ± SD; n=4. (G) IB of LdSpt16 levels in LD652 WCE 48 h post-transfection with empty vector (EV) or increasing amounts of VSV M-Flag p166 expression vector. Statistical significance in C, D, E, and F were determined by unpaired two-tailed Student’s t-test between indicated treatments. ** = P< 0.01; ns, not significant.

To directly test if LdSpt16 was involved in VSV restriction, we used *in vitro*-transcribed dsRNAs to target LdSpt16 by RNAi. Compared to control RNAi targeting an irrelevant GFP mRNA sequence, VSV-LUC gene expression (**Fig 6E**) and replication (**Fig 6F**) were significantly enhanced in LD652 cultures transfected with dsRNAs targeting LdSpt16 mRNA. These results suggested that the abortive infection of wild-type VSV in LD652 cells is at least in part dependent upon LdSpt16^SUMO^-mediated restriction. Therefore, we hypothesized that wild-type VSV M may be defective in its ability to target LdSpt16^SUMO^ for degradation. Consistent with this hypothesis, VSV M-Flag transfection into LD652 cells failed to promote LdSpt16^SUMO^ depletion at any expression level tested (**Fig 6G**). Collectively, these results suggest that VSV host range restriction in LD652 cells may be due to an inability of VSV M to antagonize LdSpt16^SUMO^ function.

### Paramyxovirus-encoded C proteins also promote proteasomal degradation of hSpt16^SUMO^

Finally, we asked if other NNS RNA viruses, unrelated to VSV, also interact with the FEAR pathway. We focused on viruses belonging to the *Paramyxoviridae* family because they include important animal and human pathogens. For example, Sendai virus (SeV) is a natural pathogen of rodents and a common cause of viral pneumonia in wild and laboratory rodents [49]. Although not known to be pathogenic to humans, SeV can productively replicate in human cells and has been an important model for understanding paramyxovirus replication and immune evasion mechanisms [50, 51].

To determine if the FEAR pathway is relevant to paramyxovirus restriction, we challenged A549 cells with a recombinant SeV strain encoding GFP (SeV-GFP) after hSpt16 RNAi. Compared to control RNAi conditions, hSpt16 RNAi led to a significant increase in SeV-GFP replication (**Fig 7A**), suggesting the FEAR pathway may also restrict paramyxoviruses. We next examined if FACT subunit levels were altered during SeV-GFP infection. Due to the slower replication cycle of SeV compared to VSV, we collected WCE at relatively later time points (8 and 24 hpi). Interestingly, while FACT subunit levels were largely unchanged at 8 hpi, hSpt16^SUMO^ subunits were specifically eliminated in SeV-GFP-infected cells by 24 hpi, suggesting SeV encodes an antagonist of hSpt16^SUMO^ (**Fig 7B**).

**Fig 7.**
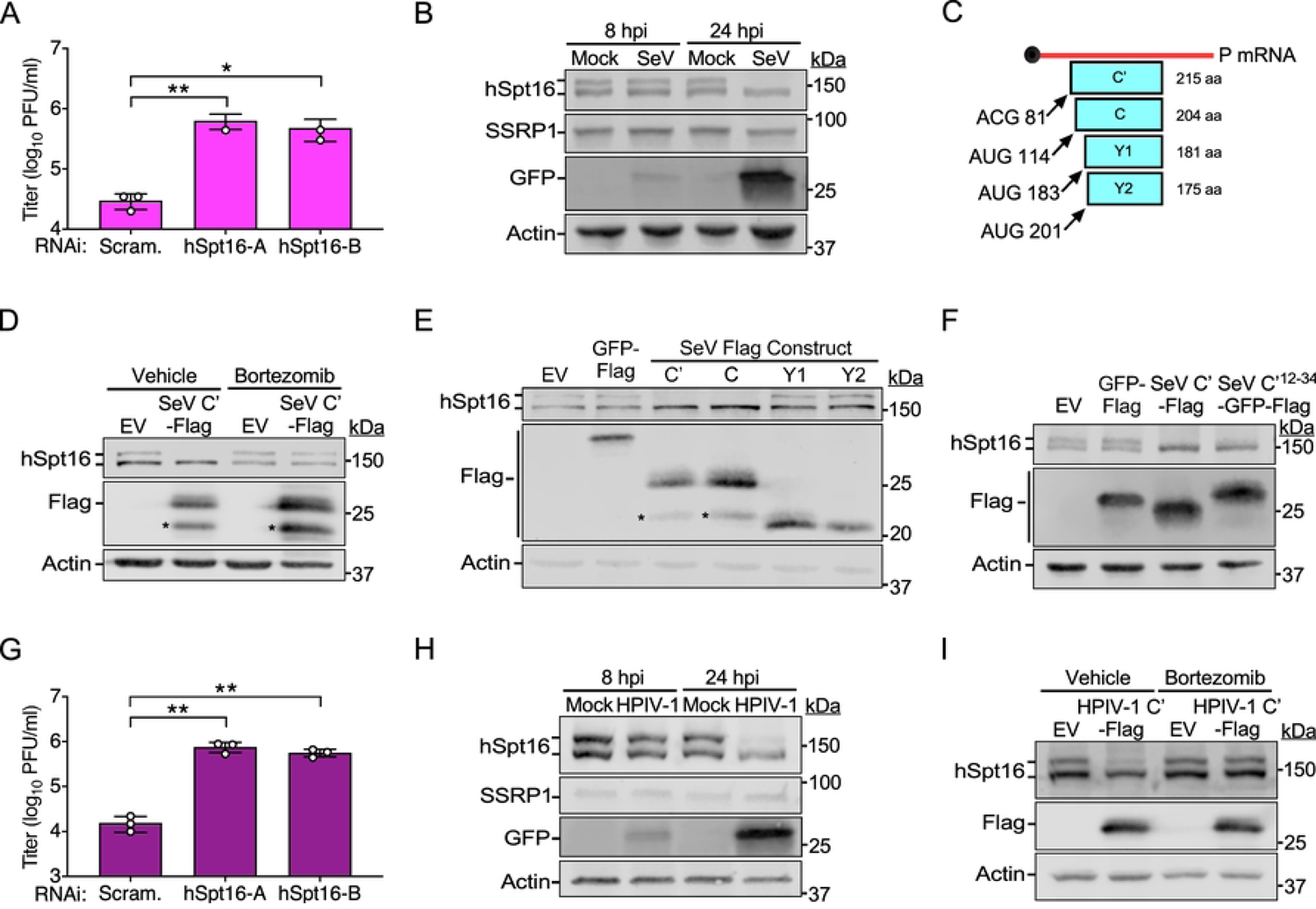
Murine and human paramyxovirus-encoded C proteins also promote proteasome-dependent degradation of hSpt16^SUMO^. (A) SeV-GFP titers 48 hpi (MOI=0.01) in A549 cells after indicated RNAi treatments. Data are means ± SD; n=3. (B) IB of FACT subunit levels in mock- or SeV-GFP-infected (MOI=1) A549 WCE at indicated hpi. GFP is a marker of infection. (C) Schematic illustrating the four accessory SeV “C” proteins (C’1, C, Y1, and Y2) and their relative translation initiation sites (nts) in the SeV *P* transcript and resulting protein sizes (a.a.). Image modified from [54]. (D) IB of endogenous hSpt16 in 293T WCE transfected with indicated expression constructs for 24 h in the absence or presence of bortezomib treatment. Asterisks indicates SeV C’-Flag degradation product. (E) IB of endogenous hSpt16 in 293T WCE transfected with indicated expression constructs for 24 h. Asterisks indicates SeV C’/C-Flag degradation products. (F) IB of endogenous hSpt16 in 293T WCE transfected with indicated expression constructs for 24 h. (G) HPIV-1-GFP titers 48 hpi (MOI=0.01) in A549 cells after indicated RNAi treatments. Data are means ± SD; n=3. (H) IB of FACT subunit levels in mock- or HPIV1-GFP-infected (MOI=1) A549 WCE at indicated hpi. GFP is a marker of infection. (I) IB of endogenous hSpt16 in 293T WCE transfected with indicated expression constructs for 24 h in the absence or presence of bortezomib treatment. Statistical significance in A and G were determined by unpaired two-tailed Student’s t-test between indicated treatments. * = P< 0.05; ** = P< 0.01; ns, not significant.

We suspected that the SeV-encoded accessory “C” proteins may be involved in mediating hSpt16^SUMO^ depletion because they function as immune evasion factors [50]. During SeV infection, C proteins are expressed as a nested set of four different proteins (C’, C, Y1, and Y2) that derive from the same SeV *P* gene-derived mRNA and contain common C-termini but differ in their N-termini due to the use of alternative translation start sites [52–54] (**Fig 7C**). SeV C’ is the largest of these four proteins so we first examined if Flag-tagged SeV C’ (SeV C’-Flag) expression could promote hSpt16^SUMO^ depletion. Interestingly, expression of SeV C’-Flag was sufficient to promote hSpt16^SUMO^ depletion in 293T cells in the absence of other SeV proteins. Moreover, this SeV C’-mediated hSpt16^SUMO^ depletion was blocked in the presence of bortezomib (**Fig 7D**). Interestingly, SeV C’-Flag could also deplete SUMOylated forms of the HA-hSpt16^ΔNLS^ mutant that cannot enter the nucleus (**Fig S6**). Importantly, SeV C’-Flag expression also reduced SUMOylated Spt16 levels in mouse 3T3 cells (**Fig S7**), indicating that this viral activity is retained in a natural SeV host species. These results collectively suggest that like VSV M, SeV C’ promotes proteasomal degradation of SUMOylated host Spt16 proteins in the cytoplasm.

We next examined if any of the shorter C proteins (C, Y1, and Y2) could also deplete hSpt16^SUMO^ levels. Interestingly, only C’ and C proteins were capable of depleting hSpt16^SUMO^ in 293T cells (**Fig 7E**), suggesting the N-terminal 23 a.a. common between C’ and C proteins, but absent in Y1 and Y2 proteins (**Fig 7C**), are critical for targeting hSpt16^SUMO^ for degradation. To determine if this 23 a.a. region was sufficient to promote hSpt16^SUMO^ depletion, we fused these residues to the N-terminus of a GFP-Flag construct, creating SeV C’^12-34^-GFP-Flag. Strikingly, expression of SeV C’^12-34^-GFP-Flag was sufficient to deplete hSpt16^SUMO^ levels (**Fig 7F**).

Human parainfluenza virus-1 (HPIV-1), also known as human respirovirus 1, is a clinically important paramyxovirus that is the leading cause of laryngotracheobronchitis (croup) in children, a disease associated with swelling of the airways and difficulty breathing [55]. HPIV-1 is closely related to SeV, sharing similar genetic and antigenic properties. For example, the C proteins of SeV and HPIV-1 are ∼70% identical [56]. Therefore, we examined the effect of hSpt16 RNAi on the replication of a GFP-encoding HPIV-1 strain (HPIV-1-GFP) and the ability of this virus to target hSpt16^SUMO^ for degradation. We found hSpt16 RNAi to enhance HPIV-1-GFP replication in human cells (**Fig 7G**). Interestingly, as with SeV infection, hSpt16^SUMO^ was specifically depleted in A549 cells by 24 hpi with HPIV-1-GFP (**Fig 7H).** Moreover, expression of a HPIV-1 C’-Flag construct in 293T cells was sufficient to deplete hSpt16^SUMO^ and this depletion was blocked by bortezomib (**Fig 7I**). Collectively, these results suggest that murine and human paramyxovirus-encoded C proteins function as FEAR pathway antagonists that target SUMOylated host Spt16 proteins for degradation.

### VSV M and paramyxovirus C proteins are structurally unrelated

Given that both VSV M and paramyxovirus C proteins promote hSpt16^SUMO^ degradation, we compared these proteins for similarity at primary (a.a. sequence) and tertiary (structural) levels. Using basic local alignment search tool (BLAST) [57] we did not observe any significant similarity between VSV M and either SeV C’ or HPIV-1 C’ sequences.

We next examined tertiary structures of these proteins. Although a full-length VSV M structure for the Indiana serotype used in our study has not been solved, the structure of a VSV M fragment encompassing the globular domain (a.a. 44-229) (**Fig S8A**) bound to Rae1-Nup98 complexes has been reported (4OWR) [58] (**Fig S8B**). In 4OWR, M contacts Rae1-Nup98 complexes using three distinct regions that project from a “fist-like” globular domain: a finger (a.a. 49-61), a thumb (a.a. 213-223), and a “web” region (a.a. 142-144). Importantly, the finger motif, encompassing the M51 residue, makes critical contacts with residues in a Rae1 β-propeller [58], illustrating the importance of this region to host interactions. Since it was our goal to compare complete VSV M and paramyxovirus C’ protein structures to one another, we generated a full-length (a.a. 1-229) VSV M structure using Alphafold2 [59](**Fig S8C**) and then compared this with 4OWR by overlaying them (**Fig S8D**) and conducting pairwise similarity tests using RSCB PDB [60] to assign root mean square deviation (RMSD) values and template modeling (TM) scores (**Fig S8E**). RMSD scores of <3 Å and TM scores >0.5 were considered to have similar overall folds while those producing RMSD values >3 Å and TM scores <0.5 were considered to be structurally unrelated [61]. Pairwise comparisons between 4OWR and our predicted VSV M structure produced a RMSD of 1.82 Å and a TM score of 0.93, providing confidence in our Alphafold2-generated VSV M structure.

We then used full-length SeV C’ and HPIV-1 C’ proteins (**Fig S9A**) to generate structures with Alphafold2. We aligned our Alphafold2 SeV C’ structure with a solved structure from a SeV C fragment (a.a. 99-204) in complex with host Alix proteins (6KP3) [62] (**Fig S9B-D**). Although 6KP3 is missing the N-terminal 23 a.a. region in SeV C’ (a.a. 12-34) involved in hSpt16^SUMO^ depletion (**Fig S9A**), it contains most of the structured globular domain (a.a. 88-215), allowing us to conduct a pairwise comparison to determine how well the Alphafold2 structure aligned with 6KP3. We obtained a RSMD value of 1.22 Å and TM score of 0.92 (**Fig S9E**), suggesting that the regions we could compare were similar in structure, giving us confidence in the Alpahfold2 model. Interestingly, the N-terminal domain of SeV C’ (a.a. 1-87) in the Alphafold2 model is completely unstructured with the exception of an alpha helix (a.a. 14-34) we term “α1” that largely encompasses the region involved in hSpt16^SUMO^ depletion (a.a. 12-34) (**Fig S9A and C**). Unfortunately, structures for HPIV-1 C proteins are not currently available, therefore we used an Alphafold2 model of full-length HPIV-1 C’ (**Fig S9F**) to overlay with our predicted SeV C’ structure (**Fig S9G**). Importantly, pairwise comparisons of these SeV and HPIV-1 structures generated a RSMD of 2.57 Å and TM score of 0.66 (**Fig S9H**), consistent with the high degree of conservation of these paramyxovirus proteins [56]. Interestingly, we again found the N-terminal region (a.a. 1-93) of HPIV-1 C’ to be entirely disordered with the exception of a short alpha helix (a.a. 18-25, α1) that corresponds to the N-terminal half of α1 in SeV C’ (**Fig S9A and G**). These data suggest that paramyxovirus C proteins have a conserved alpha helical structure that is likely important for mediating hSpt16^SUMO^ depletion.

Finally, we overlaid our Alphafold2-generated models of full length VSV M and paramyxoviruses C’ proteins (**Fig S10A**) and used RMSD and TM analyses to compare VSV M to either paramyxovirus protein. In both pairwise comparisons, we found VSV M to exhibit RMSD values >3 Å and TM scores <0.5, indicating that these proteins are structurally unrelated (**Fig S10B**). Collectively, these data suggest that VSV M and paramyxovirus proteins are structurally unrelated, supporting the notion that rhabdoviruses and paramyxoviruses evolved distinct factors to target hSpt16^SUMO^ for degradation.

## Discussion

The FEAR pathway was revealed by our discovery that poxvirus A51R proteins antagonize this pathway [6], highlighting the fact that viral immune evasion factors can provide insights into the critical host defenses that combat viral infection. While this antiviral pathway was clearly important for combating DNA viruses, its relevance to RNA viruses was less clear.

Here, we used VSV as a model RNA virus to examine if these viruses actively induce and evade the FEAR pathway. We chose this virus in part because we previously observed a measurable increase in VSV replication after hSpt16 depletion in human cells [6], hinting at the possibility that this virus may be restricted by the FEAR pathway. However, our discovery here that hSpt16 or ETS-1 depletion more strongly enhanced the replication of VSV^M51R^ versus wild-type VSV, implicated the VSV M protein as a suppressor of the FEAR pathway. Analysis of VSV-infected cell lysates led to our unexpected discovery that, unlike VV A51R, which tethers hSpt16^SUMO^ to microtubules [6], VSV M specifically promotes the proteasomal degradation of hSpt16^SUMO^ (**Fig 8**). The inability of VSV^M51R^ strains to degrade hSpt16^SUMO^ explains the strong induction of ETS-1 expression by these viruses. Importantly, the fact that VSV M expressed from a plasmid could not only deplete hSpt16^SUMO^, but also block ETS-1 induction by the ΔA51R VV strain, indicates it can antagonize ETS-1 expression independently of other VSV proteins. Moreover, our microscopy and cell fractionation studies suggest that wild-type, but not M51R mutant, VSV M also prevents ETS-1 nuclear accumulation as a second mechanism to inhibit the FEAR pathway.

**Fig 8.**
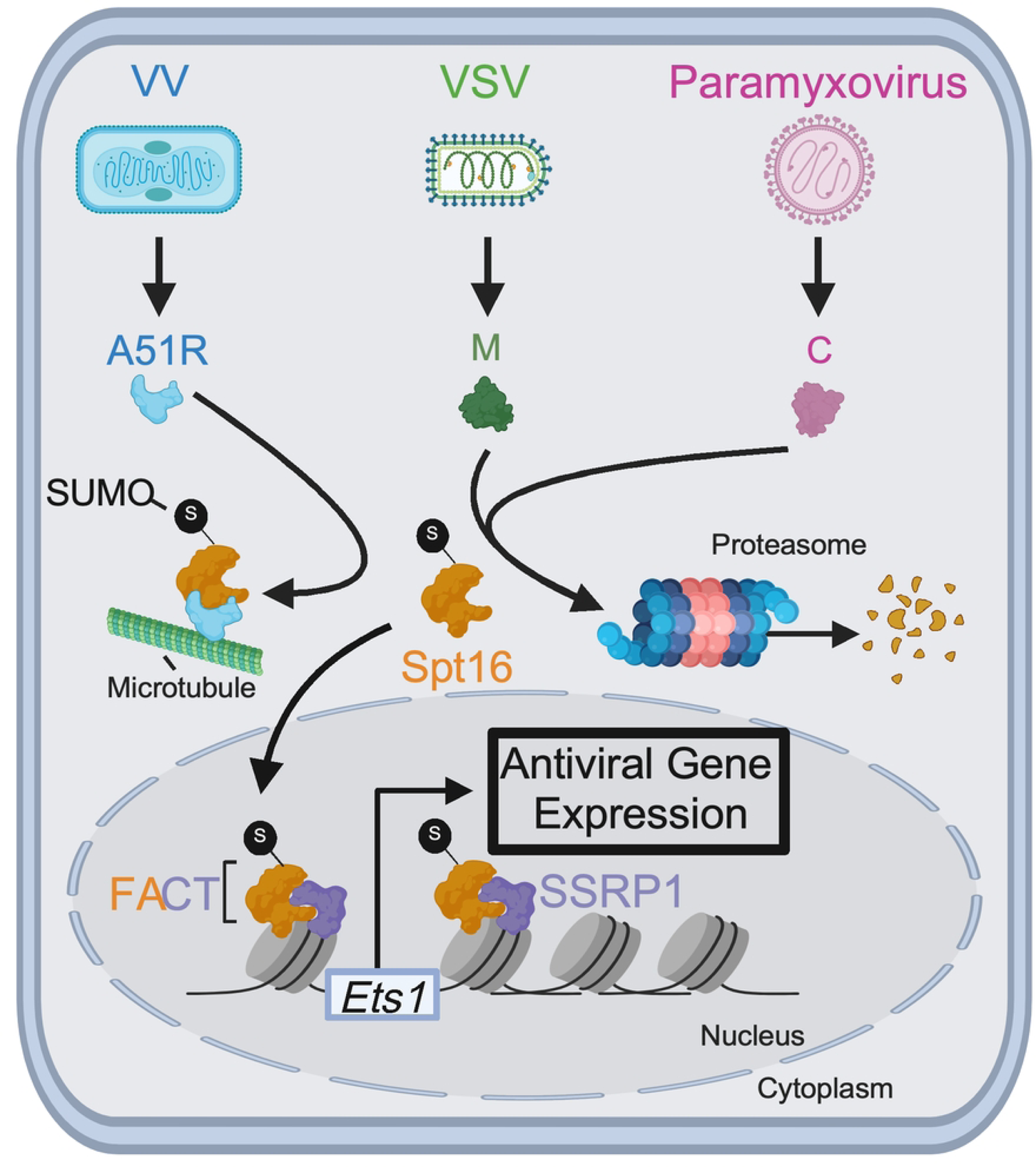
Model for DNA and RNA virus-encoded mechanisms to antagonize the FEAR pathway. Poxvirus A51R proteins tether hSpt16^SUMO^ subunits to microtubules to prevent FACT complex formation and activation of ETS-1 expression in the nucleus. In contrast, VSV M and paramyxovirus (SeV, HPIV-1) C proteins promote degradation of hSpt16^SUMO^ subunits in the proteasome to block ETS-1 expression. This degradation-based mechanism may involve recruitment of host E3 ubiquitin ligases to target hSpt16^SUMO^ for degradation. VSV M may also inhibit FEAR pathway activation by impeding ETS-1 nuclear accumulation (not shown). Model for FEAR pathway is adapted from [6] and was created using Biorender.com.

The molecular mechanisms by which VSV M promotes hSpt16^SUMO^ degradation and prevents ETS-1 nuclear accumulation are active areas of investigation. Given that the most well-characterized defect of the M51R M mutant is its inability to interact with the host Rae1-Nup98 mRNA export complex, we were surprised that Rae1 or Nup98 depletion had no effect on VSV-mediated hSpt16^SUMO^ depletion. Given that the M51 residue resides in a “finger-like” projection that protrudes from the globular domain of VSV M [58], it may be a docking site for not only Rae1 but also for additional host factors. We speculate that VSV M may use this finger projection to recruit ubiquitin-proteasome system-related machinery (e.g. E3 ubiquitin ligases) that target hSpt16^SUMO^ for degradation (**Fig 8**). Indeed, the repurposing of host E3 ligases to degrade antiviral host factors has been widely observed among DNA and RNA viruses [63, 64]. Prior work has shown that VSV M can inhibit nuclear translocation mediated by specific karyopherin/importin protein family members in xenopus oocytes and this activity is lost in the M51R mutant [65, 66]. Thus, it is possible that VSV M interferes with import factors required for ETS-1 nuclear translocation.

Our discovery that VSV^ΔM51^/ VSV^M51R^ strains are defective in FEAR pathway antagonism has led to important implications for their use in oncolytic virotherapy. A major drawback of these strains is their inability to effectively replicate in cancer cell types that are still capable of mounting effective antiviral responses [17, 40, 67]. Prior work had shown that 786-0 cells are highly refractory to VSV^ΔM51^ but that compounds termed “viral sensitizers” that inhibit IFN responses can enhance the replication in, and killing of, these cells by this oncolytic strain [18, 21, 68]. Our data demonstrating that hSpt16 RNAi enhances the replication of a VSV^ΔM51^strain in these cells suggest that IFN-independent responses (i.e. the FEAR pathway) are also relevant to the restriction of this strain. Moreover, we found the FACT inhibitor, CBL0137, to be a promising viral sensitizer that improved VSV^ΔM51^ replication in these and other human cancer lines known to be resistant to infection by VSV^ΔM51^/VSV^M51R^ oncolytic strains. Importantly, CBL0137 is already being pursued in clinical trials as a cancer chemotherapeutic [33] as FACT upregulation is often observed during oncogenesis and is associated with more aggressive cancers [31]. Such FACT upregulation may also lead to more potent FACT-dependent antiviral responses, potentially posing barriers to oncolytic virus replication in tumors. Thus, it will be important to determine in future studies if combined (VSV^ΔM51^+CBL0137) treatment can improve oncolytic virotherapy efficacy in animal models.

This study has also provided insight into the poorly understood abortive infection of VSV in lepidopteran insect cells and the rescue of VSV replication in these cells by poxvirus A51R proteins [9]. We previously found Spt16 proteins from various mammalian, insect, and nematode cell lysates to exhibit the same, two-band pattern on IBs found in human cell extracts, suggesting SUMOylation of Spt16 proteins is highly conserved among both vertebrate and invertebrate eukaryotes [6]. More formal evidence for this was provided here where we demonstrated that treatment with a SUMOylation inhibitor (tannic acid) [47] could specifically eliminate the higher molecular weight form of LdSpt16 in *L. dispar*-derived LD652 cells. Strikingly, despite being encoded by a mammalian poxvirus, VV A51R was able to interact with LdSpt16^SUMO^. This is likely due to the high degree of conservation of the A51R-binding domain [6] among human and moth Spt16 proteins. Our findings that: 1) VSV M cannot deplete LdSpt16^SUMO^ levels; 2) A51R proteins lacking a functional Spt16-binding domain cannot rescue VSV replication in *L. dispar* cells; and 3) LdSpt16 RNAi rescues VSV replication, all strongly implicate LdSpt16^SUMO^ in VSV host range restriction in these cells. Importantly, these results also suggest an ancient role for SUMOylated Spt16 proteins in eukaryotic antiviral immunity. Future studies will be needed to determine if insect ETS transcription factors [69] are also involved in virus restriction and whether the FEAR pathway is indeed intact in invertebrates.

Investigation of paramyxovirus-FEAR pathway interactions revealed a remarkable example of independent evolution of a common immune evasion mechanism by NNS RNA viruses. Despite a lack of primary a.a. sequence or structural homology between VSV M and paramyxovirus C proteins, they could both promote proteasome-dependent degradation of hSpt16^SUMO^ (**Fig 8**). The specific targeting of SUMOylated Spt16 proteins by pox-, rhabdo-, and paramyxoviruses suggests these host factors activate a critical node of antiviral immunity. Interestingly, SeV and HPIV-1 C proteins have also been shown to antagonize STAT protein function to inhibit IFN signaling [50, 51]. Thus, SeV and HPIV-1 are likely under similar evolutionary pressure to inhibit key immune responses pathway components such as STAT1 and Spt16^SUMO^ in their murine and human hosts. Future studies will be aimed at determining if paramyxovirus C and VSV M use a common or distinct host interactions to ultimately achieve hSpt16^SUMO^ degradation, but our current work clearly defines these proteins as the first RNA virus-encoded FEAR pathway antagonists.

In conclusion, our work has demonstrated that RNA viruses can activate and evade the FEAR pathway, indicating this pathway is relevant to this important class of pathogens. Major questions remain regarding how RNA viruses activate the FEAR pathway and how ETS-1 activation eventually leads to virus restriction. Studies are ongoing to explore these exciting areas of FEAR pathway biology. Moreover, whether other RNA virus families antagonize the FEAR pathway is an active area of investigation. Ultimately, a greater understanding of the activation and evasion of the FEAR pathway by viruses will help us better appreciate the role of this host response in the evolutionary arms race between viruses and their hosts.

## Materials and Methods

Specific details regarding the source of all key experimental reagents (primers, plasmids, Abs, viruses, cell lines, etc.) can be found in **Table S1**.

### Cell lines and primary cultures

Mammalian cell lines were maintained at 37°C in 5% CO_2_ atmosphere. A549, U2OS, 293T, 786-0, NIH 3T3, Vero, and BHK-21 cells were cultured in DMEM supplemented with 10% FBS. RPMI 7951 cells were cultured in EMEM supplemented with 10% FBS. BSC-40 cells were passaged in MEM containing 5% FBS. All media above additionally contained 1% non-essential amino acids, 1% L-glutamine, and 1% antibiotic/antimycotic (Gibco). LD652 were cultured in a 1:1 mixture of Ex-Cell 420 (Sigma) and Graces insect medium (Invitrogen) supplemented with 10% FBS at 27°C under normal atmospheric conditions [9, 70].

### Viruses and virus titrations

VSV-GFP and VSV^ΔM51^-GFP strains were amplified in either Vero or BHK-21 cells using low MOI infections [9, 71]. All other VSV strains were amplified using low MOI infections in BHK-21 cells. SeV-GFP and HPIV-1-GFP stocks were obtained from Viratree. BSC-40 cells were used for VV stock preparation [9]. Unless otherwise noted, virus strains were titrated by plaque assay (VSV-LUC, ΔA51R^FA51R^, ΔA51R) or fluorescent foci/plaque assay (VSV-eGFP, VSV^M51R^-eGFP, VSV^ΔM51^-GFP, SeV-GFP, HPIV-1-GFP) on BSC-40 cell monolayers, with a 1.5% low-melting point agarose (Invitrogen) overlay used for VSV titrations [6]. In some cases, VSV-GFP and VSV^ΔM51^-GFP stocks were titered on Vero cells [71]. Experimental viral infections were incubated for 1 h in serum free DMEM at 37°C before the inoculum was replaced with complete media for the remainder of the infection with the exception of SeV-GFP and HPIV-1 cultures, which were kept in serum free DMEM containing either 1% TrypLE Select (Invitrogen) or 5 μg/ml acetylated trypsin (Sigma). Where indicated, replacement media after 1 h of infection contained indicated doses of CBL0137 (Selleck Chemicals) and was kept in the media until the end of the experiment. At indicated times post-infection, infected cell culture supernatants were collected, clarified by centrifugation, and clarified supernatant titers were determined by serial dilution followed by plaque assay or fluorescent foci/plaque assay [6].

### Expression vectors

The human ETS-1-HA pEZ-M07 expression vector was obtained from GeneCopoeia. VSV M-Flag and M51R/ΔM51 mutant derivatives were constructed by gene synthesis (Gene Universal) with inclusion of *Sac*II and *Pac*I sites for cloning into *Sac*II/*Pac*I-digested pcDNA3 [6]. VSV M-Flag was also cloned into the insect p166 expression vector [9] using *Sac*II/*Pac*I sites. SeV C’-Flag and HPIV-1 C’-Flag expression constructs were also generated using gene synthesis and cloned into pCNDA3 using flanking *Sac*II/*Pac*I sites. SeV C-, Y1-, and Y2-Flag expression plasmids were constructed from PCR amplification from the full-length SeV C’ sequence. The SeV C’^M12-R34^-GFP-Flag pcDNA3 construct was made from PCR amplifying GFP-Flag from pcDNA3 with a forward primer encoding the M12-R34 sequence. All SeV-related constructs were cloned into pCDNA3 using flanking *Sac*II/*Pac*I sites. Wild-type VV Flag-A51R and Flag-A51R^158-162Ala^ open reading frames were cloned into the *Sac*II/*Pac*I sites of p166 by *Sac*II/*Pac*I digest of Flag-A51R and Flag-A51R^158-162Ala^ pcDNA3 vectors [6]. The HA-hSpt16 and HA-hSpt16^ΔNLS^ pEZ-M06 expression vectors have been described [6]. The Flag-GFP pcDNA3 and p166 vectors have been described [6, 9]. All genes were sequence verified. Expression plasmids were transfected into indicated mammalian cell lines as previously described using Lipofectamine 2000 (Invitrogen) in Opti-MEM I (Gibco) [9] overnight and then media was replaced with complete culture media. Plasmids transfected into LD652 cells were used FuGENE HD (Promega) (LD652 cells) [72]. Cells were for the indicated times prior to further processing [e.g. protein extraction (see below)].

### General protein extraction and inhibitor treatments

Cells were washed with phosphate-buffered saline (PBS) prior to scraping and transfer into a 1.5 mL microcentrifuge tube for centrifugation at 800 x *g* at 4°C for 15 min. Cell pellets were resuspended in either 1x Reporter Lysis Buffer (Promega) containing cOmplete™ EDTA-free Protease Inhibitor Cocktail (Roche) and 1 mM phenylmethylsulfonyl fluoride (PMSF) and freeze-thawed prior to addition of 5x SDS-PAGE loading buffer (100 mM tris HCl, pH 6.8, 4% SDS, 12% (v/v) glycerol, 4 mM DTT, 0.02% (w/v) Bromophenol Blue) or were resuspended in Pierce RIPA buffer (containing cOmplete™ EDTA-free Protease Inhibitor Cocktail and 1 mM PMSF) prior to addition of 5x SDS-PAGE loading buffer. Where indicated, cells were treated with 10 µM tannic acid for 4 h prior to protein harvest. Also, where indicated, bortezomib (Selleck Chemicals) (25 nM), MG132 (Sigma) (40 μM), or DMSO (Corning) (vehicle control) was added to cells 6 h post-transfection and was kept in media until protein extraction.

### Immunoblotting

WCE were boiled for 10 min prior to SDS-PAGE electrophoresis at 50 V for approximately 4 h. Separated proteins were transferred in Towbin Buffer (BioRad) onto nitrocellulose membranes at 150 mA at 4°C for 90 min and blocked with Odyssey Blocking Buffer (LI-COR) for 1 h at RT. Membranes were blotted with primary Ab overnight at 4°C, with actin serving as a loading control. After 3 x 5 min PBS-T (PBS, 0.1% Tween) washes, membranes were incubated in secondary Ab conjugated to an IRDye (LI-COR) for 1 h, washed 3 x 5 min in PBS-T, then a final 5 min PBS wash. Membranes were then imaged with an Odyssey Fc Imager (LI-COR).

### Immunoprecipitation

Infection-based co-immunoprecipitation experiments were performed as described with minor modifications [6]. Briefly, 10^6^ LD652 cells were seeded overnight, then infected with ΔA51R^FA51R^ (MOI=25) for 24 h. Prior to cell lysis, cells were washed with equal volumes of PBS twice. Cells were lysed in IP lysis buffer (cOmplete™ EDTA-free Protease Inhibitor Cocktail, 1 mM PMSF, 25 mM Tris-HCl at pH 7.4, 150 mM NaCl, 0.5% NP-40) and subjected to shearing and sonication (two-15 second sonications with a 30 second interval). Samples were benzonase (Sigma) treated (250 units/mL) for 1 h at RT. 10% of “input” was collected, and remaining lysate was end-over-end incubated with 5 µg of primary Ab overnight at 4°C (rabbit-anti-Flag or isotype control Ab). Then, lysates were incubated with PureProteome protein A/G magnetic beads (Sigma) for 1-2 h, extensively washed, and immunoprecipitants eluted in 60 μl 2x SDS-PAGE loading buffer. IPs of total SUMOylated protein fractions in mock-, VSV-eGFP, or VSV^M51R^-eGFP-infected (MOI=10) A549 cells for 12 h used either SUMO-1 or SUMO-2/3 Ab included with the Signal Seeker SUMOylation 1 or 2/3 Detection Kit (Cytoskeleton) and IPs were carried out according to the manufacturer instructions [6].

### Cell fractionation

Cell fractionation and subsequent densitometric analyses were conducted as previously described [6]. Briefly, cells were infected at the indicated MOI and times for the indicated times prior to fractionation using the Subcellular Protein Fractionation Kit (Thermo Scientific) into either cytosolic or nuclear fractions. Densitometry-based band quantifications were performed using ImageJ software (NIH, version 1.51n). It is important to note that because the lysis buffer volume used in the fractionation procedure is dependent upon the initial cell pellet size (which varied between infection conditions), and the fractionation buffers were not compatible with protein quantification assays, we only compared cytosolic versus nuclear distribution of a particular protein within (and not between) each infection condition [6]. Quantification of the percentage of total cellular ETS-1 protein present in the nuclear fraction within each indicated infection condition was calculated by dividing the nuclear fraction ETS-1 band densitometric values by the sum of total (cytosolic +nuclear) ETS-1 densitometric values.

### RNAi in cell culture

All siRNAs were obtained from Sigma’s pre-designed siRNA library (see **Table S1)**. The hSpt16 and ETS-1-targeted siRNAs have been previously validated [6]. Transient siRNA-mediated knockdown was achieved by reverse transfecting mammalian cells with Lipofectamine 2000 according to manufacturer’s protocol for 48-72 h prior to viral infection [6]. RNAi knockdowns in LD652 cells were performed as previously described using *in vitro*-transcribed dsRNAs transfected with Cellfectin II (Invitrogen) in Sf-900 II serum free medium [9]. Transfection medium was replaced with complete culture medium after overnight culture. Sequences of primers used to generate dsRNA targeting GFP (control) or *LdSpt16* gene sequences are in **Table S1**.

### Luciferase assays

Luciferase assays in LD652 WCE were performed as previously described with minor modifications [9, 72]. Briefly, at the indicated times post-infection, cells were washed in PBS, collected by centrifugation (2000 rpm, 10 min, 4°C) and lysed in reporter lysis buffer (Promega). Lysates were spotted to 96-well dishes, mixed with Luciferase Assay Reagent (Promega) and arbitrary light units (LU) were measured using an FLUOstar Omega Plate reader (BMG Labtech).

### Antibodies

Information regarding antibodies used in this study, including their source, is available in **Table S1**.

### Fluorescence microscopy and cell viability assays

At indicated times post-infection, cells were stained with 200 μL of serum free media containing NucBlue (Thermo Fisher) for 30 min followed by replacement with 1 mL of PBS. Cells were imaged using a 4X objective on an EVOS-FL fluorescence microscope (Thermo Fisher) using DAPI and GFP cubes. Each condition had three replicate wells, and at least four images/well were collected for analysis. Image analysis was conducted using Fiji (NIH) to quantify the normalized GFP signal/field by dividing total GFP signal by total DAPI (NucBlue) signals. Positive signal was determined by using uninfected wells lacking NucBlue stain to set a minimum threshold in both DAPI and GFP channels that was applied to the remaining dataset. Fold change in GFP signals were calculated by dividing normalized GFP values of experimental treatments with normalized GFP values in control treatments (indicated in each figure). For GFP foci imaging and cell viability analysis of 786-0 cells infected with VSV^ΔM51^-GFP, cells were grown to confluency in multi-well dishes, infected with VSV^ΔM51^-GFP at the indicated MOIs and treated with 0.3-3µM CBL0137 (or media alone) 1 hpi using the Agilent Bravo liquid handling system (Agilent, CA, USA). 24 hpi, GFP images were captured by fluorescence microscopy using an ArrayScan Thermo Scientific Cellomics) and GFP foci were quantified. Cell viability was assessed 48 hpi using resazurin sodium salt (Sigma-Aldrich) after a 2.5 h incubation using a BioTek Microplate Reader (BioTek Instruments Inc.).

### Immunofluorescence microscopy

For staining cells under mock, VSV-eGFP, or VSV^M51R^-eGFP infection conditions, cells were seeded at a density of 30,000 cells per coverslip, cultured overnight, then infected (MOI=10) for 12 hpi. Cells were fixed with 4% paraformaldehyde (Sigma), incubated with blocking buffer (PBS with 1% BSA and 0.1% Triton-X) for 1 h, stained with mouse anti-ETS-1 Ab for 2 h, washed extensively with blocking buffer, incubated with Alexa Fluor-conjugated secondary Ab for 1h, and then again washed extensively. Coverslips were then mounted onto glass slides using ProLong™ Diamond anti-fade with DAPI (Thermo Scientific) and imaged with an Olympus FV10i confocal laser scanning microscope (version 2.1) equipped with Olympus Fluoview software (version 4.2a). Images were deconvoluted with CellSens Imaging software (version 1.18). Similar imaging methods were used for U2OS cells transiently transfected with VSV M-Flag, VSV M^M51R^-Flag and ETS-1-HA expression plasmids, except rabbit anti-Flag and mouse anti-HA primary Ab were used for staining.

### Protein sequence alignment and structure prediction

Sequences of VSV M (NCBI accession: 7UMK_B), SeV C’ (P04862.3), and HPIV-1 C’ (NP_604434.1) were predicted using AlphaFoldv2 [59] and analyzed using PyMOL (The PyMOL Molecular Graphics System version 1.2r3pre, Schrödinger, LLC.). To determine structural similarity between VSV M, SeV C’ and HPIV-1 C’, pairwise structural alignments were conducted using RSCB PDB [60] to assign RMSD and TM scores. RMSD values of <3 Å and TM scores >0.5 were considered to have similar overall fold, while pairwise comparisons producing RMSD values >3 Å and TM scores <0.5 were considered to be structurally unrelated [61].

### Statistical analyses

All statistical analyses were conducted using GraphPad Prism (version 8.0) software and *P* values <0.05 were considered statistically significant. Sample sizes, statistical tests used, and *P* value information are indicated in the respective figure or figure legend for each quantitative experiment.

## Data Availability

The authors declare that the main data supporting the findings of this study are available within the article and/or Supplementary Information. Source data will be provided after initial peer review as per the editor’s instructions as Supplementary Information files due to the size and time required to prepare.

## Author Contributions

Conceptualization, E.A., R.A., J.S.D., and D.B.G.; Methodology, E.A., R.A., J.S.D., and D.B.G.; Investigation, E.A., D.S., A.E., R.A., M.M.S., and D.B.G.; Data Curation, E.A., D.S, A.E., R.A., M.M.S., and D.B.G.; Writing—original draft, E.A. and D.B.G.; Writing—Review & Editing, E.A., D.S., A.E., R.A., M.M.S., J.S.D., and D.B.G.; Visualization, E.A., A.E., R.A., and D.B.G. Supervision, D.B.G., R.A., and J.S.D.; Funding Acquisition, J.S.D and D.B.G.

## Acknowledgments

We thank members of the Gammon lab for helpful discussion regarding the manuscript. We also thank Dr. Beatriz Fontoura (UTSW Medical Center) for advice regarding the manuscript. This work was supported by NIH grants 1R35GM137978 and 1R21AI180551, and American Cancer Society Research Scholar grant RSG-23-1153296-01-DMC to DBG. EAR and AE were supported by NIH Training Grant T32 AI007520.

## Abbreviations

FEAR: FACT-ETS-1 Antiviral Response
VSV: vesicular stomatitis virus
M: matrix
IFN: interferon
FACT: facilitates chromatin transcription
VV: vaccinia virus
Spt16: suppressor of Ty 16 homologue
hSpt16: human Spt16
hSpt16^SUMO^: SUMOylated hSpt16
SSRP1: structure specific recognition protein 1
ETS-1: E26 transformation-specific sequence-1
RanGAP: Ran GTPase activating protein
Rae1: ribonucleic acid export 1
Nup98: nucleoporin 98
NNS: nonsegmented negative-strand
LdSpt16: *Lymantria dispar* Spt16
LdSpt16^SUMO^: SUMOylated LdSpt16
LUC: firefly luciferase
SeV: Sendai virus
HPIV-1: Human parainfluenza virus-1.

## Supplementary Figure Legends

**Fig S1.**
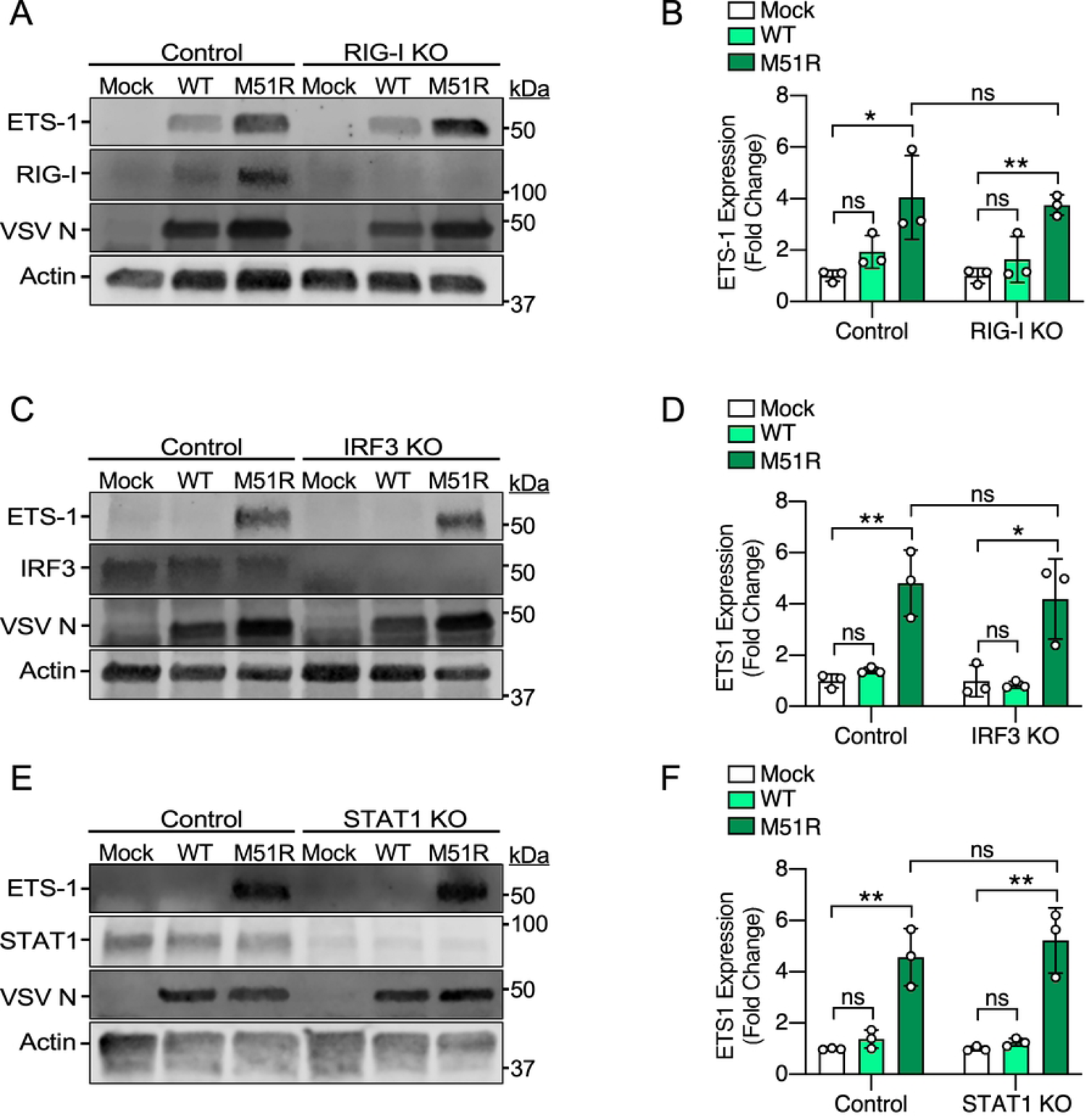
VSV-induced ETS-1 expression is independent of RIG-I or IFN signaling. (A, B) Representative IB (A) and quantification (B) of ETS-1 expression using IB of WCE from control or RIG-I knockout (KO) A549 cells infected with the indicated strains (MOI=10) for 8 h. Note: RIG-I is expressed at a low level in uninfected cells and is known to be induced by viral infection [73, 74], consistent with our results. (C, D) Representative IB (C) and quantification (D) of ETS-1 expression using IB of WCE from control or IRF3 KO A549 cells infected with the indicated strains (MOI=10) for 8 h. (E, F) Representative IB (E) and quantification (F) of ETS-1 expression using IB of WCE from control or STAT1 KO A549 cells infected with the indicated strains (MOI=10) for 8 h. Data in B, D, and F are means ± SD for n=3 experiments. Statistical significance in B, E, and F were determined by unpaired two-tailed Student’s t-test between indicated treatments. For brevity, only the most relevant statistical comparisons are shown. * = P< 0.05; ** = P< 0.01; ns, not significant.

**Fig S2.**
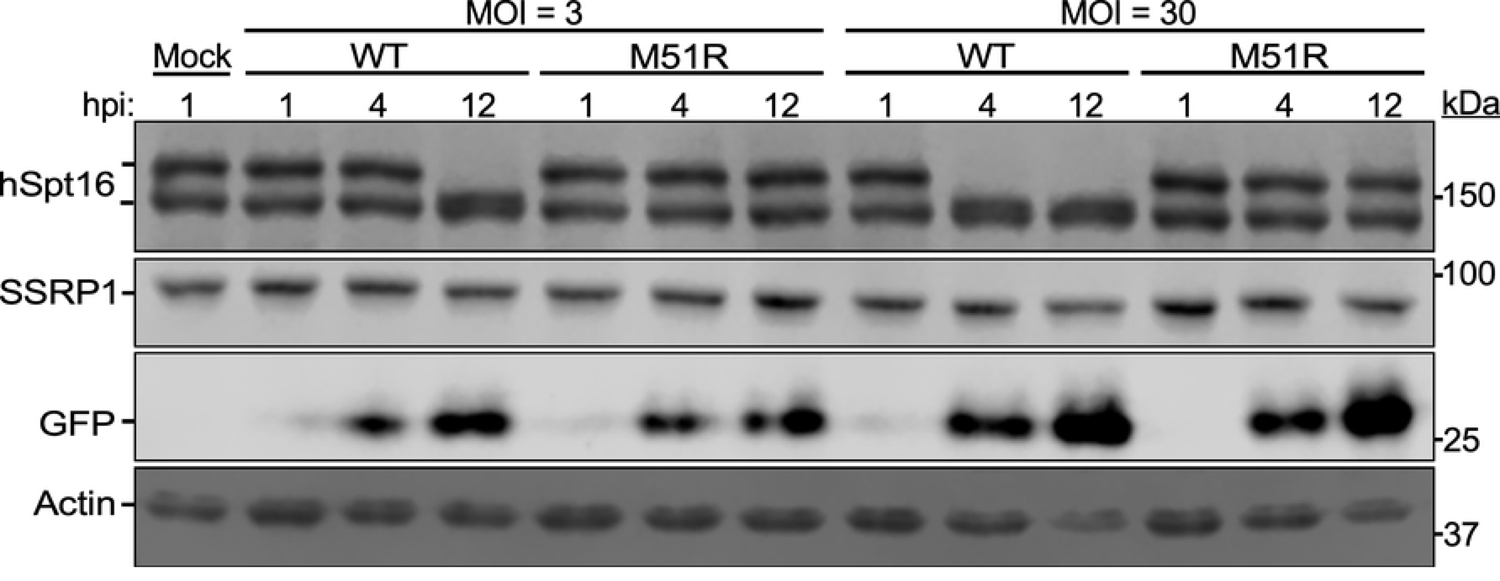
The VSV^M51R^-eGFP strain is unable to deplete hSpt16^SUMO^ regardless of MOI. IB of endogenous hSpt16 in A549 WCE after infection with VSV-eGFP or VSV^M51R^-eGFP at the indicated MOI. GFP is used as a marker for infection.

**Fig S3.**
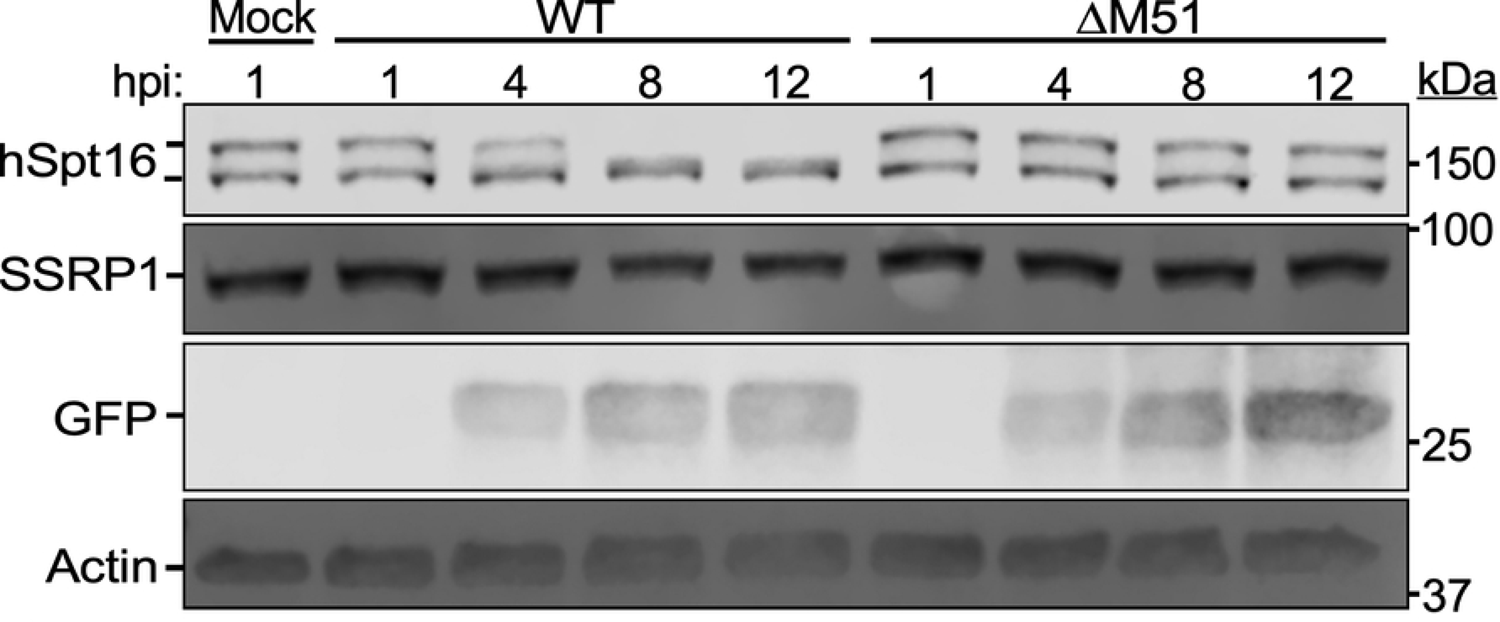
A VSV^ΔM51^ strain is unable to deplete hSpt16^SUMO^. IB of endogenous hSpt16 in A549 WCE after infection with VSV-GFP (WT) or VSV^ΔM51^-GFP (ΔM51) (MOI=3). GFP is used as a marker for infection.

**Fig S4.**
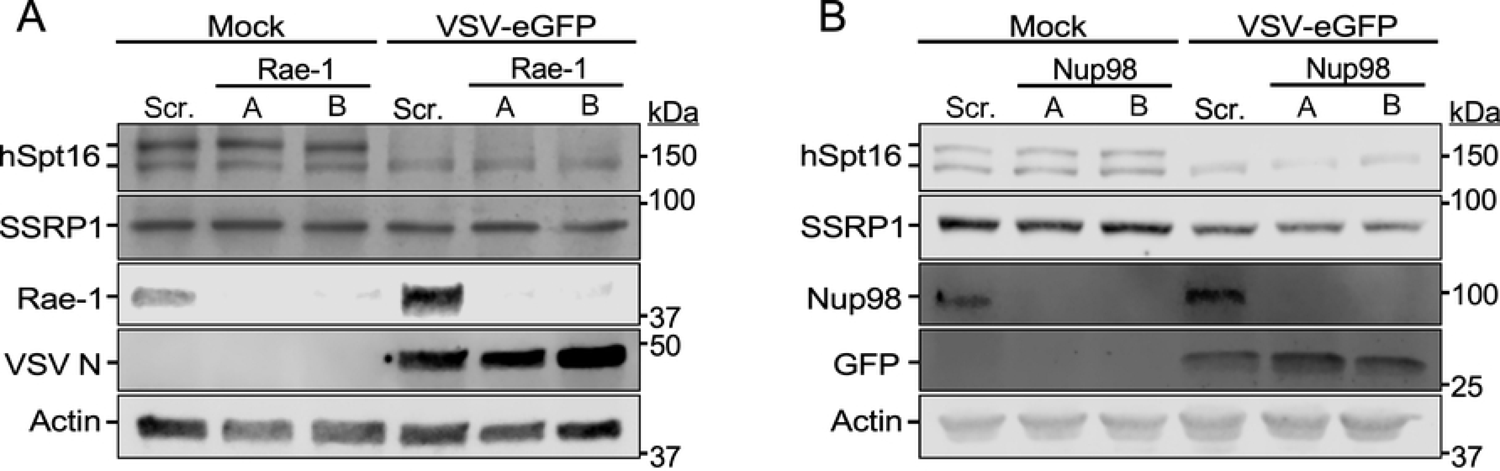
VSV M-mediated depletion of hSpt16^SUMO^ is independent of Rae1 and Nup98. IB of endogenous hSpt16 in A549 whole cell extract (WCE) 72 h after RNAi of Rae1 (A) or Nup98 (B) under mock- or VSV-eGFP-infection (MOI = 10) conditions for 12 h. VSV N and GFP are markers for infection. Scram., scrambled.

**Fig S5.**
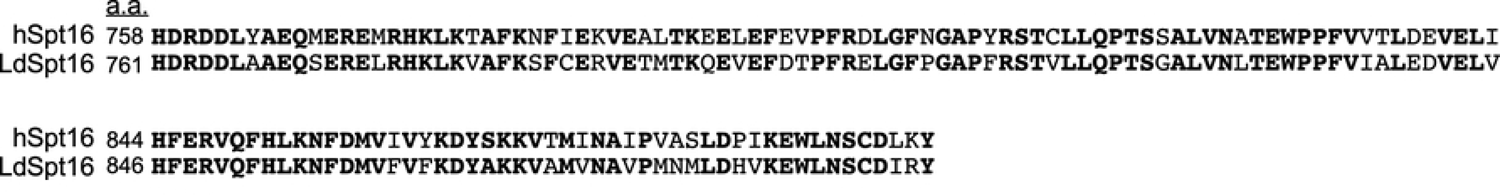
Human and *Lymantria dispar*-encoded Spt16 proteins share a similar A51R-binding domain. Alignment of the VV A51R-binding domain of hSpt16 [6] (accession: NP_009123.1) with corresponding region in LdSpt16 (accession: GCA_004115105.1). The overall amino acid identity of these regions is 72.8%. Alignment was made using NCBI Blast2p Software.

**Fig S6.**
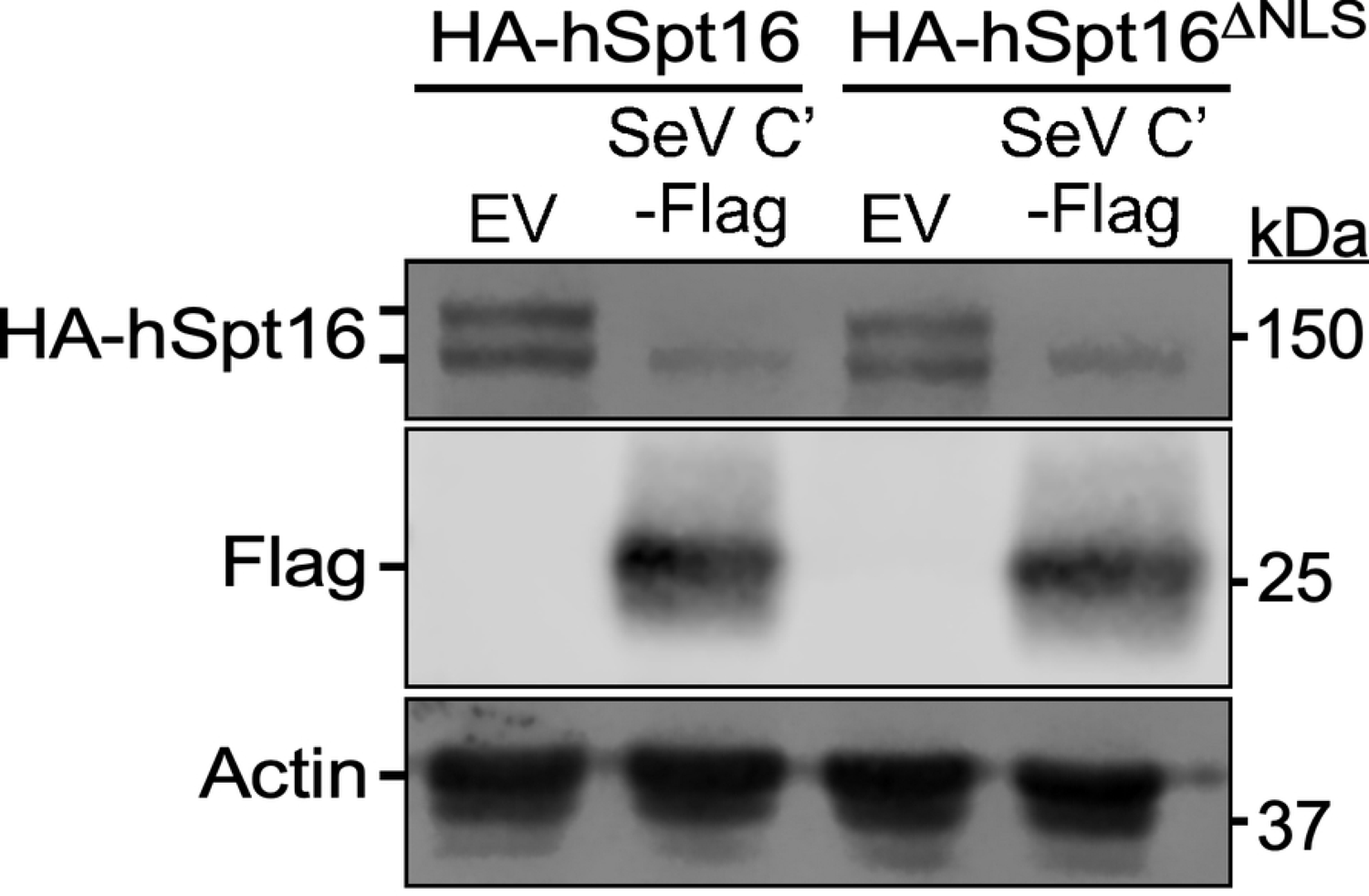
SeV C’ promotes depletion of SUMOylated Spt16 subunits present in the cytosol. IB of HA-hSpt16 or HA-hSpt16^ΔNLS^ in 293T WCE 24 h after transfection with empty vector (EV) and or SeV C’-Flag expression constructs.

**Fig S7.**
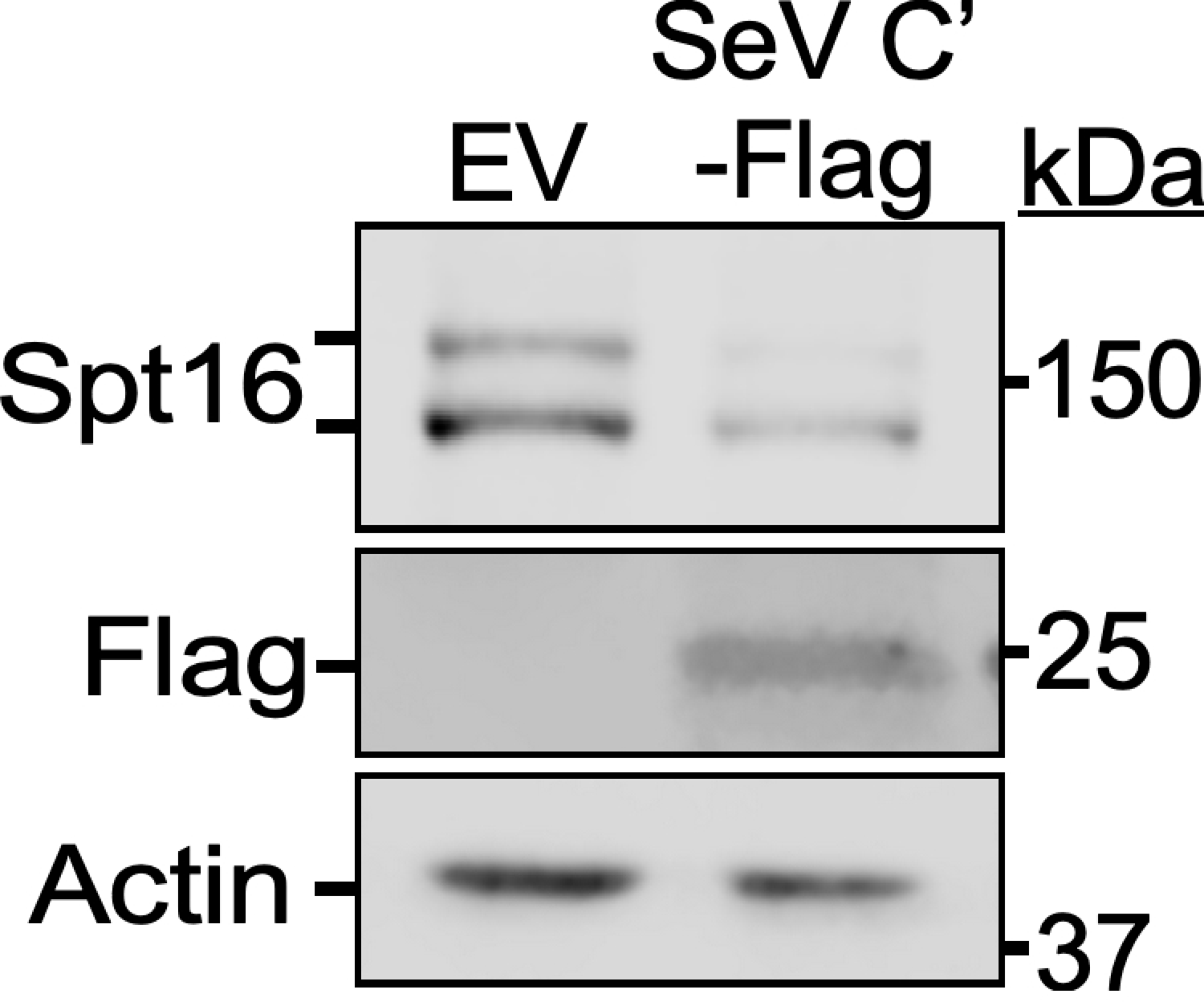
SeV C’ promotes depletion of SUMOylated Spt16 subunits in mouse 3T3 cells. IB of endogenous mouse Spt16 in 3T3 WCE 24 h after transfection with empty vector (EV) and or SeV C’-Flag expression constructs.

**Fig S8.**
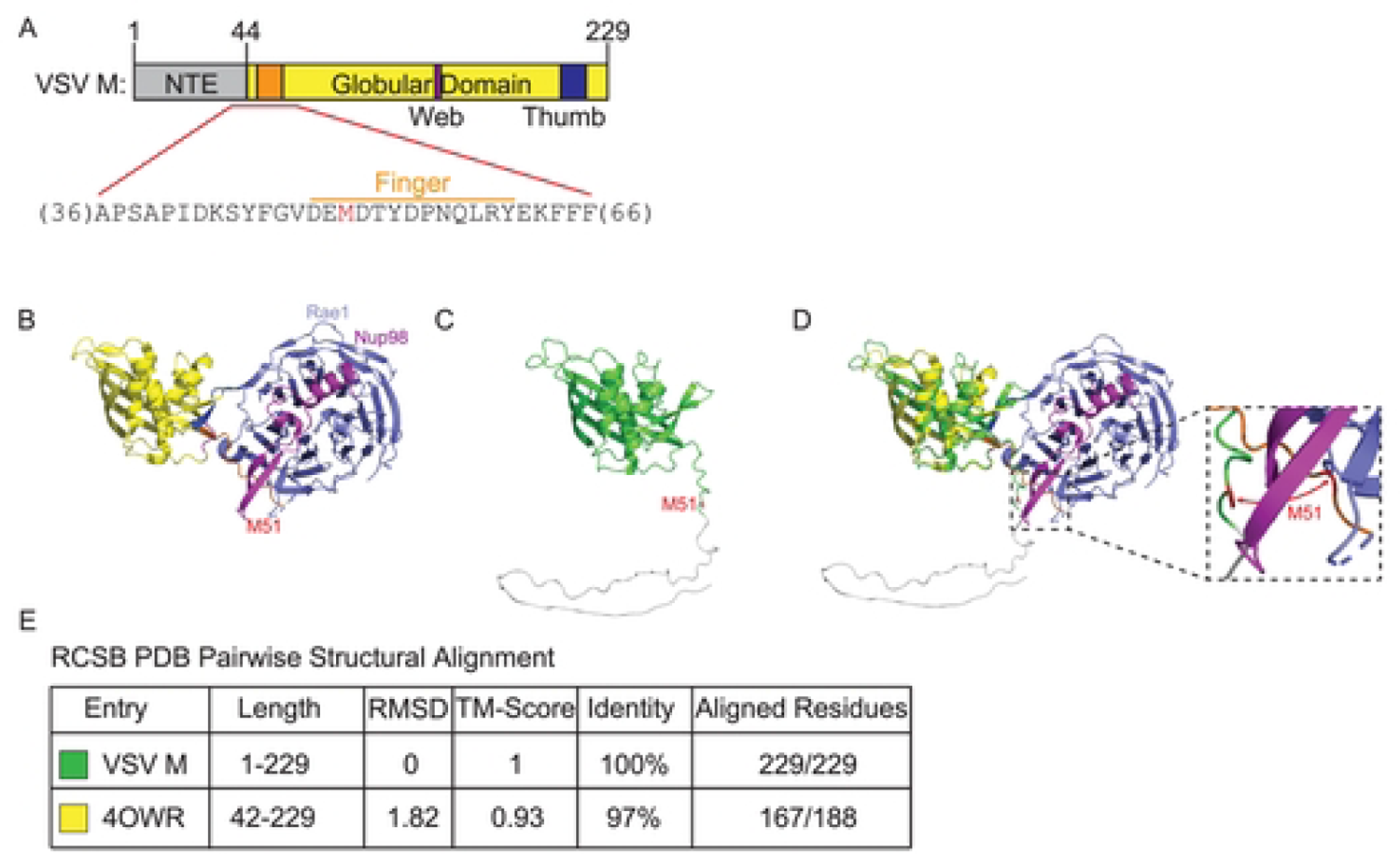
VSV M protein structural modeling. (A) VSV M protein map showing region surrounding M51 residue (red). NTE = N-terminal extension. Protein map based on reference [58]. B) 4OWR structure of VSV M fragment (a.a. 44-229) bound to Rae1-Nup98 complex [58]. (C) Alphafold2-generated structure of full-length (a.a. 1-229) VSV M (strain Indiana). (D) Overlay of structures from B and C. E) Results of pairwise comparison of structures in B and C showing RMSD (in Å) and TM values.

**Fig S9.**
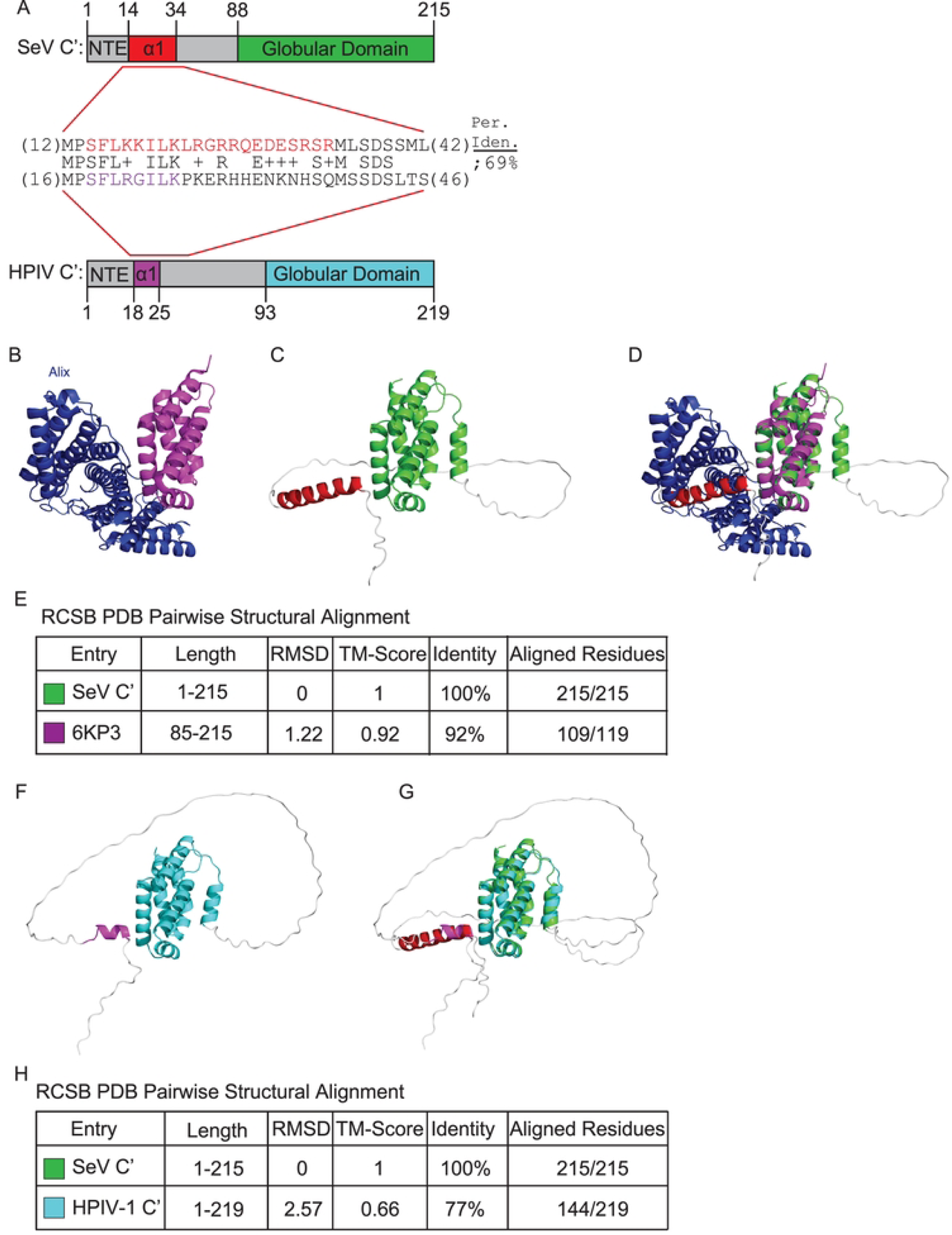
Paramyxovirus C protein structural modeling. (A) Protein maps of SeV C’ and HPIV-1 C’ showing region in SeV C’ N-terminus sufficient for hSpt16^SUMO^ degradation (a.a. 12-34) aligned with corresponding region in HPIV-1 C’. NTE = N-terminal extension. Residues predicted to form alpha helical structures in the NTE are in red and purple for SeV C’ and HPIV-1 C’, respectively. (B) 6KP3 structure of SeV C’ fragment (a.a. 99-204) bound to cellular Alix protein [62]. (C) Alphafold2-generated structure of full-length (a.a. 1-215) SeV C’. (D) Overlay of structures from B and C. (E) Results of pairwise comparison of structures in B and C showing RMSD (in Å) and TM values. (F) Alphafold2-generated structure of full-length (a.a. 1-219) HPIV-1 C’. (G) Overlay of structures from C and F. (H) Results of pairwise comparison of structures in B and C showing RMSD (in Å) and TM values.

**Fig S10.**
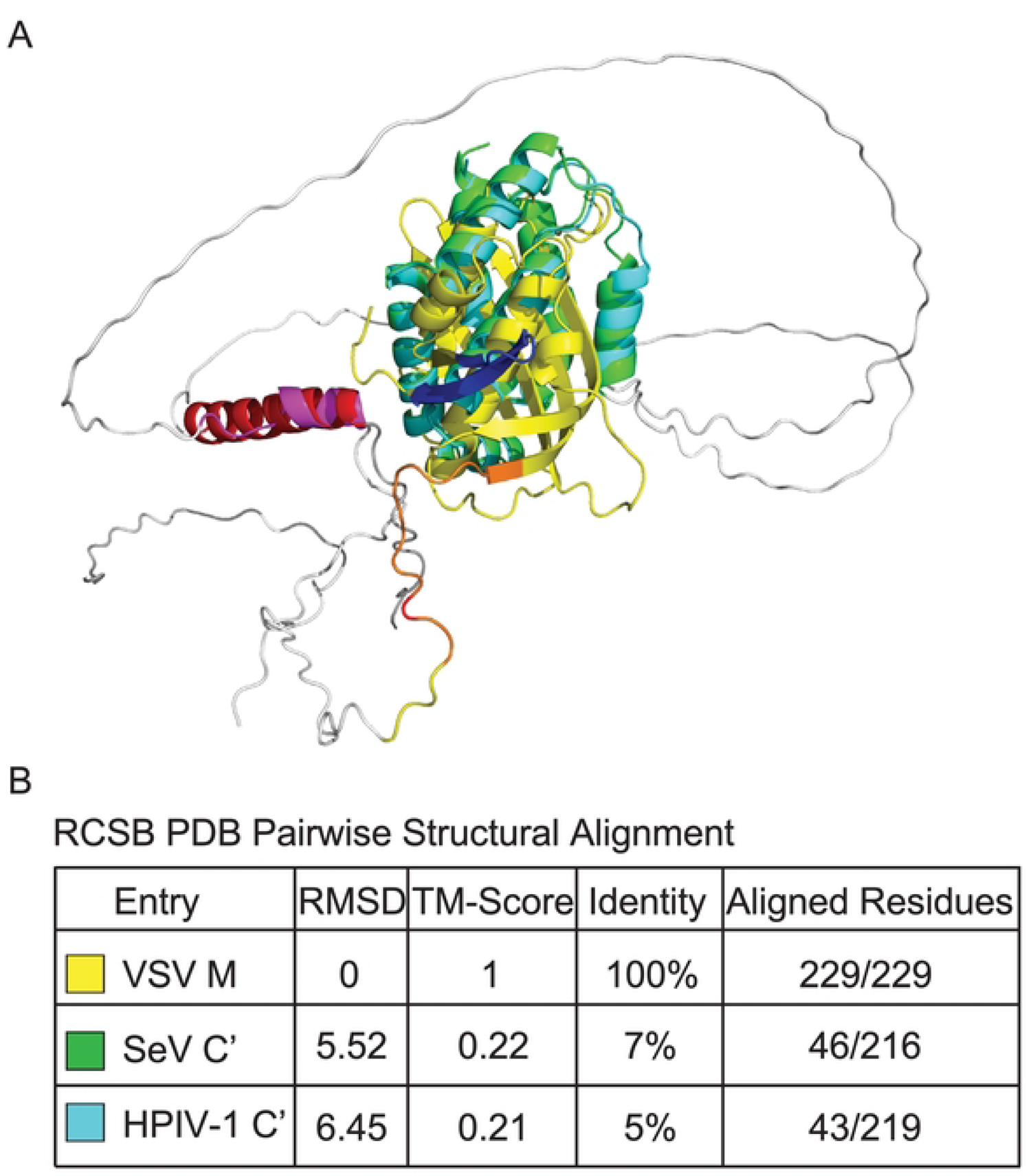
Structural comparison of VSV M to paramyxovirus C proteins. (A) Overlay of Alphafold2-generated VSV M (Fig S8C), SeV C’ (Fig S9C), and HPIV-1 C’ (Fig S9F) structures. (B) Results of pairwise comparison of VSV M to either SeV C’ or HPIV-1 C’ structures showing RMSD (in Å) and TM values.

## Supplementary Tables

**Table S1.** Key Experimental Reagents.

